# Identification and characterization of dietary antigens in oral tolerance

**DOI:** 10.1101/2024.05.26.593976

**Authors:** Jamie E. Blum, Ryan Kong, E.A. Schulman, Francis M. Chen, Rabi Upadhyay, Gabriela Romero-Meza, Dan R. Littman, Michael A. Fischbach, Kazuki Nagashima, Elizabeth S. Sattely

## Abstract

Food antigens elicit immune tolerance through the action of intestinal regulatory T (Treg) cells. Unlike food allergens, the proteins that mediate tolerance are mostly undescribed. Here, we found that epitopes derived from seed storage proteins are targets of murine intestinal Treg cells, with the most frequent response targeting the C-terminus of the maize protein alpha-zein. An MHC tetramer loaded with this antigen revealed that zein-specific T cells are predominantly intestinal Treg cells, develop concurrently with weaning, and constitute up to 2% of the peripheral Treg cell pool. Zein-responsive Treg cells repressed naïve T cell proliferation *ex vivo* and prior dietary exposure resulted in a constrained response upon multiple inflammatory challenges in vivo, supporting a specific role for gut-resident Treg cells in suppressing systemic immune responses. Together, our work reveals the development, immune suppressive characteristics, and function of naturally occurring Treg cells that recognize dietary seed-storage proteins, a previously undescribed class of antigens in oral tolerance.

**One-sentence summary:** Immunodominant epitopes from seed storage proteins are a target of intestinal regulatory T cells that modulate immune challenge to food.

## INTRODUCTION

Humans consume nearly 100 g of protein a day from varied sources. Despite being non-self, these foods typically result in oral tolerance, a phenomenon defined by a state of immune unresponsiveness following subsequent exposure to a given antigen (*1, 2*). Oral tolerance is an intrinsic function of the immune system resulting from continuous surveillance of intestinal contents. Orchestration of oral tolerance and immune suppression is thought to be mediated by regulatory T (Treg) cells, a subset of CD4 T cells that recognize distinct dietary epitopes through a unique T cell receptor (TCR) (*3, 4*). Prior work has pointed to multiple mechanisms for Treg cell-mediated immune suppression, including anti-inflammatory cytokine secretion (e.g., interleukin (IL)-10, TGFβ), competition for growth factors (e.g., IL-2 binding by CD25), and cell-cell inhibition (e.g., Lag3, LAP, CTLA4). These mediators can suppress conventional T cell activation directly or via reduced antigen presentation (*5*).

The induction of durable and specific immune tolerance is critical for allergy prevention, and its restoration is the primary goal of allergy immunotherapy. Despite the interest in programming antigen-specific Treg cell responses, few dietary proteins have been identified that mediate tolerance, and none in an untargeted manner that reveals naturally selected epitopes (*6*). Repertoire level analyses have shown that antigen-free diets change the abundance and composition of intestinal T cells (*3, 7*). In one report, nearly half of the peripherally induced Treg cells emerge when mice are exposed to dietary protein, and thus presumably recognize a food epitope, though specific antigens were not identified (*3*). Most studies of oral tolerance use model antigens in adoptive transfer paradigms – these antigens differ in dose, route, and timing of exposure, possibly eliciting transient phenotypes; and focusing on an adoptively transferred cell occludes the epitope selection step, which is integral to initiating an immune response (*8*– *12*). Recently, in studies to identify immune epitopes from gut resident bacteria, our team serendipitously identified TCRs responsive to a component of mouse chow (*13*). Here, we describe the identification and characterization of multiple dietary antigen-TCR pairs and phenotype these food-responsive T cells in vivo to reveal an in-depth profile of the molecular events that result in an oral tolerance response.

## RESULTS

### Identification of dietary antigens from corn, soy, and wheat

In order to identify the food epitopes recognized by T cells, we adopted a strategy recently developed for the discovery of T cell epitopes from the gut microbiome. In this prior work, mice were colonized with a complex defined bacterial community, intestinal T cells were isolated, and single cell-sequencing was used to identify TCRs (*13*). We then constructed T cell hybridomas bearing these receptors and performed an *in vitro* stimulation assay, incubating them individually with each bacterium in the community. We observed four T cell hybridomas that were unresponsive to any of the bacterial strains but were strongly induced by germ-free stool or chow (*13*), suggesting the possibility that the corresponding TCRs respond to a food-derived epitope. Notably, each of these TCRs were found predominantly (or exclusively) on Treg cells, suggesting they mediate a tolerogenic response. These data are consistent with our, and previously reported, repertoire level analyses of mice showing that consumption of protein-containing chow increases the population of gut resident Treg cells compared to amino acid defined (AAD) diets (fig. S1) (*3*).

In this work, we set out to determine which specific components in chow are recognized by Treg cells as a critical step toward characterizing the molecular events that lead to durable oral tolerance to food encountered through the gut. We first screened TCRs found on sequenced Treg cells, selected using a variety of strategies (Table S1), for chow responsiveness. In total 128 hybridoma cell lines each bearing a unique TCR were generated and screened. To identify the antigens recognized by these TCRs, we began by incubating the T cell hybridomas with homogenized preparations of each of the 7 protein-containing components of mouse chow (wheat, corn, oat, fish meal, soybean, alfalfa, and yeast), using dendritic cells for antigen presentation (Fig. 1A-B). Five of the TCRs were activated by corn, while the other two were specific for soy and wheat. Food-responsive TCRs were cloned from Treg cells obtained from both germ-free and colonized animals, together resulting in a total set of seven food-responsive TCRs

**Fig. 1:**
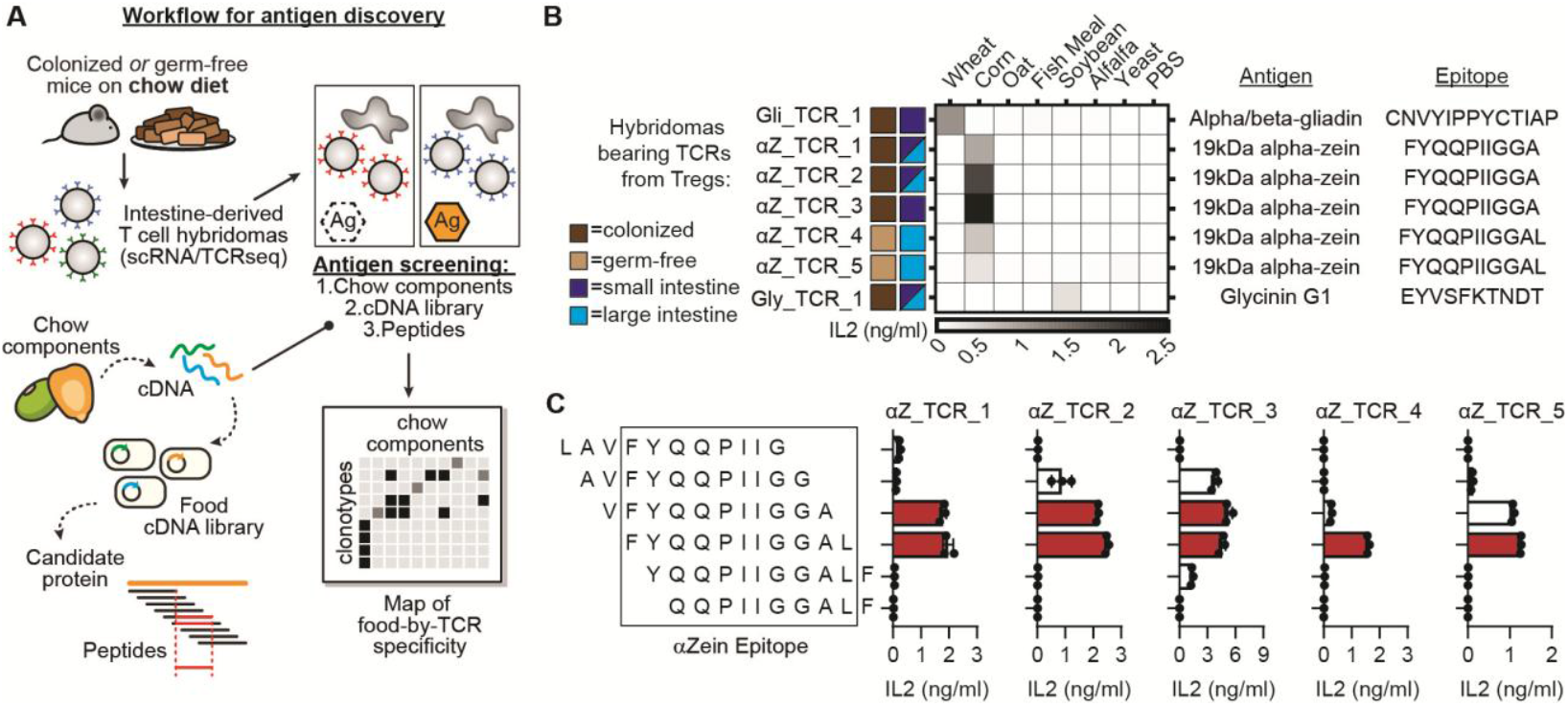
Epitope-TCR pairs identified from wheat, corn, and soy. **(A)** Workflow to screen TCR hybridomas to find food-responsive TCRs from single cell sequencing data, map the cognate antigens using a cDNA library screen, and map exact epitopes using synthetic peptides. **(B)** TCRs were selected as described in Table S1. A mixed lymphocyte assay was used to determine TCR-responsiveness to dietary components, antigens and epitopes. **(C)** Tiling scans with 10-11 amino acid peptides spanning alpha-zein were used to identify the minimum antigen epitope that was recognized by each TCR as indicated by IL-2 secretion. Red bars do not differ significantly from each other (p > 0.05) and represent the maximum activation. N=3 replicates/condition. *P* values were calculated using a one-way ANOVA with Tukey’s multiple comparisons test to make all possible pairwise comparisons. Error bars indicate mean ± standard deviation. Every dot represents a cell culture replicate. TCR, T cell receptor; DC, dendritic cell.

We next used an untargeted approach to identify the cognate antigen of each TCR, starting with the five corn-responders. We expressed a corn cDNA library in *E. coli* (Fig. 1A), with the anticipation this library would approximate the seed proteome (estimated 10,000 proteins (*14*)). Approximately 17,280 total clones were selected and initially tested in 576 bins of 30. *E. coli* cells were heat-killed and the resulting lysates were tested for their ability to activate food-responsive hybridomas. Bins that stimulated a response were retested as individual clones, and those yielding signal were sequenced. All five of the corn-specific TCRs were restimulated by an isoform of 19 kDa alpha-zein, a member of a family of closely related maize proteins that differ in sequence and size (*15*) (TCRs henceforth called αZ_TCRs). We next tested a set of tiled synthetic peptides to identify the epitope recognized from within the protein (Fig. 1A). Strikingly, we found that the C-terminal sequence FYQQPIIGGAL (αZein_223-233_) is the epitope for all 5 αZ_TCRs (Fig. 1B-C; Fig. 2A). In addition to dendritic cells, αZ_TCRs were effectively stimulated by CX3CR1^+^ macrophages, which are known to facilitate antigen sampling in the intestine (fig S2A, (*16*)). To our knowledge, this epitope has not previously been associated with a T cell response in mice or humans. The αZein_223-233_-responsive TCRs were no more similar in sequence to each other than to other food responsive TCRs (fig. S2B). The sole exception - αZ_TCR_1 and αZ_TCR_5 – differed by only 1 amino acid. However, they derived from different mice, suggesting independent convergence on a nearly-identical TCR sequence. Taken together, these data suggest that αZein_223-233_ is an immunodominant epitope.

**Fig. 2:**
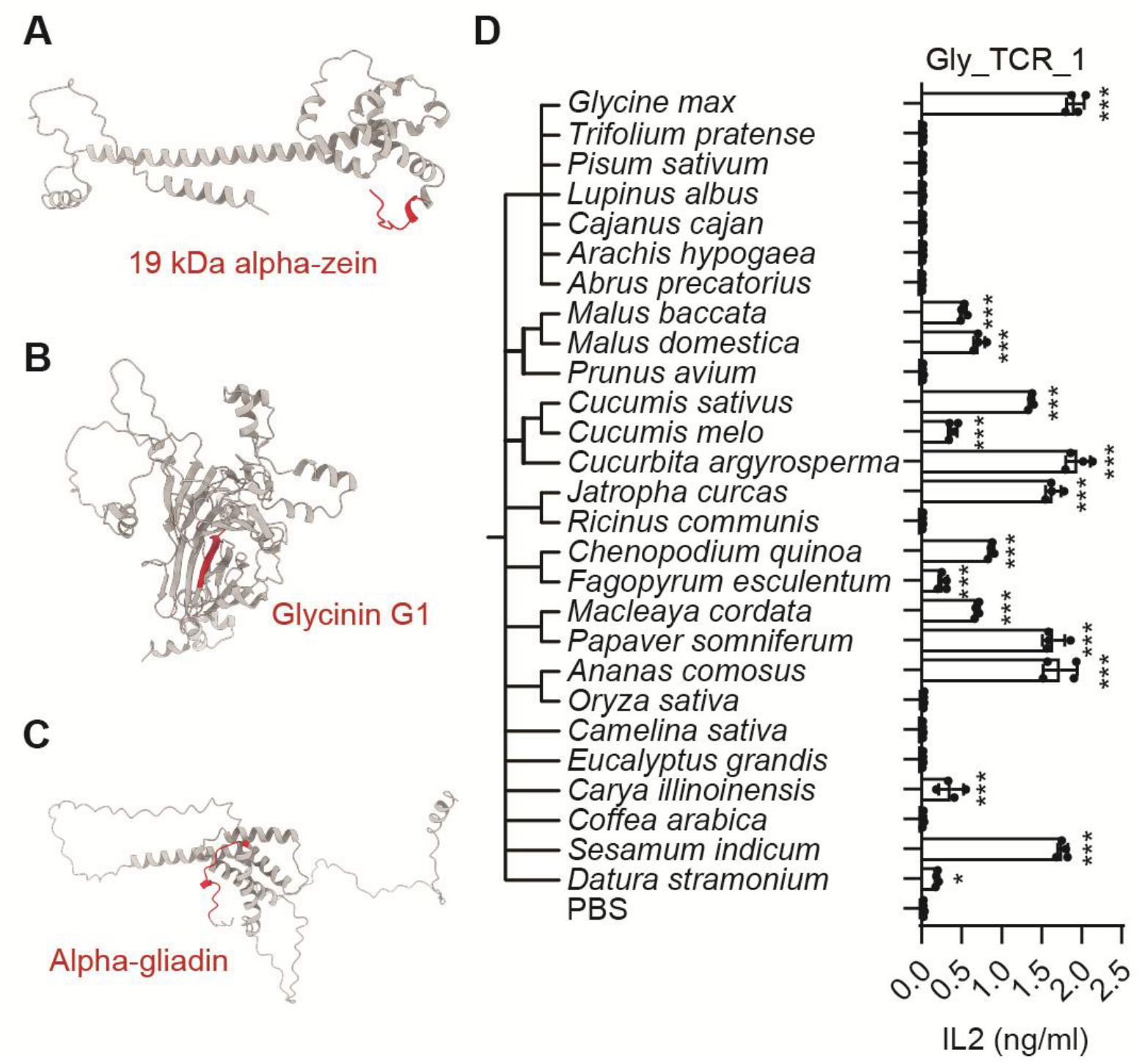
Identified epitopes from alpha-zein, Glycinin G1 and gliadin and cross-reactivity of Gly_TCR_1. **(A-C)** Alpha-fold predicted structure of alpha-zein, glycinin G1, and gliadin with identified epitopes highlighted in red. **(D)** Gly_TCR_1 was stimulated in the hybridoma mixed lymphocyte assay with soybean or lysates from seeds containing a putative soybean homolog. N=4/condition. *P* values were calculated using a one-way ANOVA and Dunnett’s multiple comparisons test was used to compare each lysate against the PBS control. Error bars indicate mean ± SD. Every dot represents a cell culture replicate. * p<0.05, ** p<0.01 and *** p<0.001 compared to the PBS control.

Seed storage proteins were also the targets of the soy and wheat-responsive TCRs. We identified the epitope EYVSFKTNDT from the soy protein glycinin G1 (Glycinin_430-439_) using a screening approach similar to that described for corn. The soy epitope resides partially in an internal beta sheet of this protein (Fig. 1B; Fig. 2B; fig S2C). This epitope is orthogonal to known Glycinin G1 epitopes, which were previously described as antibody binding regions from soy-allergic patients(*17*). Further, we found the wheat-responsive TCR recognizes the recently-reported epitope CNVYIPPYCTIAP (Gliadin_273-285_) (Fig. 1B, fig. S2D)(*6*). In prior work in B6 mice, this epitope was found to be a target of the T cell response to intraperitoneal injection of gliadin, suggesting some degree of peptide-intrinsic dominance across different immune contexts. Like αZein, gliadin belongs to the water-insoluble prolamin group of seed storage proteins and the epitope is situated within a C-terminal region that is predicted to be unstructured (Fig. 2C).

### Conservation of TCR epitopes across the plant kingdom

Immune cross-reactivity has been well documented between allergens (*18*–*22*), but it is unknown whether tolerogenic epitopes also display cross-reactivity. Notably, the gliadin_273-285_ epitope is directly conserved across numerous wheat/grass species and lysates from these species activated Gli_TCR_1 (fig. S2E, data file S1). A computational search for homologous peptides revealed orthologs of the epitope recognized by the soy-specific TCR in the genomes of many other plants. To test for cross-reactivity, we incubated the Gly_TCR_1 T cell hybridoma with seed lysates from 26 plants, along with dendritic cells. We found that Gly_TCR_1 is robustly activated by seed lysates from numerous plants (Fig. 2D); notable elicitors included other foods such as *Carya illinoinensis* [pecan], *Chenopodium quinoa* [quinoa], and *Sesamum indicum* [sesame seed]. Glycinin is an 11S globulin protein, and homologous 11S globulins could explain the cross-reactive signal in other species (data file S2, e.g., 11S globulin seed storage protein 1-like from quinoa with the sequence EWVSFKTND). Orthologs of this epitope are also conserved in foods we did not assess (e.g. walnut, pistachio, guava) and in trees (e.g., cork oak, white poplar). Curiously, plants that are more closely related to soybean – e.g., *Arachis hypogaea* [peanut], *Pisum sativum* [pea]) – were less cross-reactive than more distantly related species.

While Gly_TCR_1 recognized closely related variants of the soy epitope Glycinin_430-439_, the αZ_TCRs were more specific. Synthetic peptides that represent other isoforms of 19 kDa αZein with similar sequences elicited no response in the same assay (fig. S3A). Seed lysates from *Panicum miliaceum* and *Setaria italica*, which contained predicted αZein_223-233_ homologs, mildly activated only αZ_TCR_3 (fig. S3B, data file S3). It is possible that αZein_223-233_ homologs from other plants are expressed at a low level, different developmental stage, or only under particular environmental/stress conditions. Since processing and presentation of possible epitopes by antigen-presenting cells (APCs) was not assessed, weak or lack of signal in this assay does not exclude potential protein cross-reactivity. To directly test activation in response to a homolog, αZ_TCRs were stimulated with equal quantities of a synthetic peptide from a *Panicum miliaceum* homolog or the cognate zein epitope. While these results highlighted the potential for cross-reactivity, they support a preference for αZein_223-233_ (fig. S3C). Thus, taken together, some epitopes are directly conserved (e.g., gliadin), some have close cross-reactive homologs (e.g., glycinin), and some have close homologs but exhibit little cross-reactivity (e.g., zein).

### αZein-specific T cells are predominantly small intestine Treg cells

The T cell epitopes we identified are recognized in the physiological setting of a natural food matrix encountered by ingestion—rather than, e.g., a purified protein in drinking water or a food-derived extract administered by injection. Given that the TCRs that recognize these epitopes were predominantly identified on Treg cells, we reasoned that αZein _223-233_-, glycinin_430-439_-, and gliadin_273-285_-responsive T cells may mediate oral tolerance to food.

To characterize the distribution and functional properties of αZein_223-233_-responsive T cells, we obtained an MHCII tetramer loaded with αZein_223-233_, the corn-derived epitope recognized by five of the seven TCRs we mapped. We first analyzed the context in which αZein-specific Treg cells emerge. In young adult (6-12 wk old) mice fed a standard chow diet, small intestine αZein_223-233_ tetramer-positive T cells were predominantly Foxp3^+^ Treg cells, and a majority expressed the transcription factor RORγt across mice from 2 different vendors (Fig. 3A-B, fig. S4A). αZein-specific T cells were also detected in the large intestine (Fig. 3C). Compared to the small intestine, fewer numbers of αZein-specific T cells were detected in large intestine precluding definitive analysis of cell type distribution, although some Treg cells were observed (fig. S4B). On average, greater than 2% of small intestine pTreg (Foxp3^+^Helios^-^ or Foxp3^+^RORγt^+^) cells were tetramer-positive, a strikingly large response against a single food epitope (fig. S4C). This result suggests the possibility that a subset of food epitopes dominate immune recognition in the context of tolerance. αZein_223-233_-specific T cells were also detected in the mesenteric lymph nodes, although to a much smaller degree than intestine (fig. S4D).

**Fig. 3:**
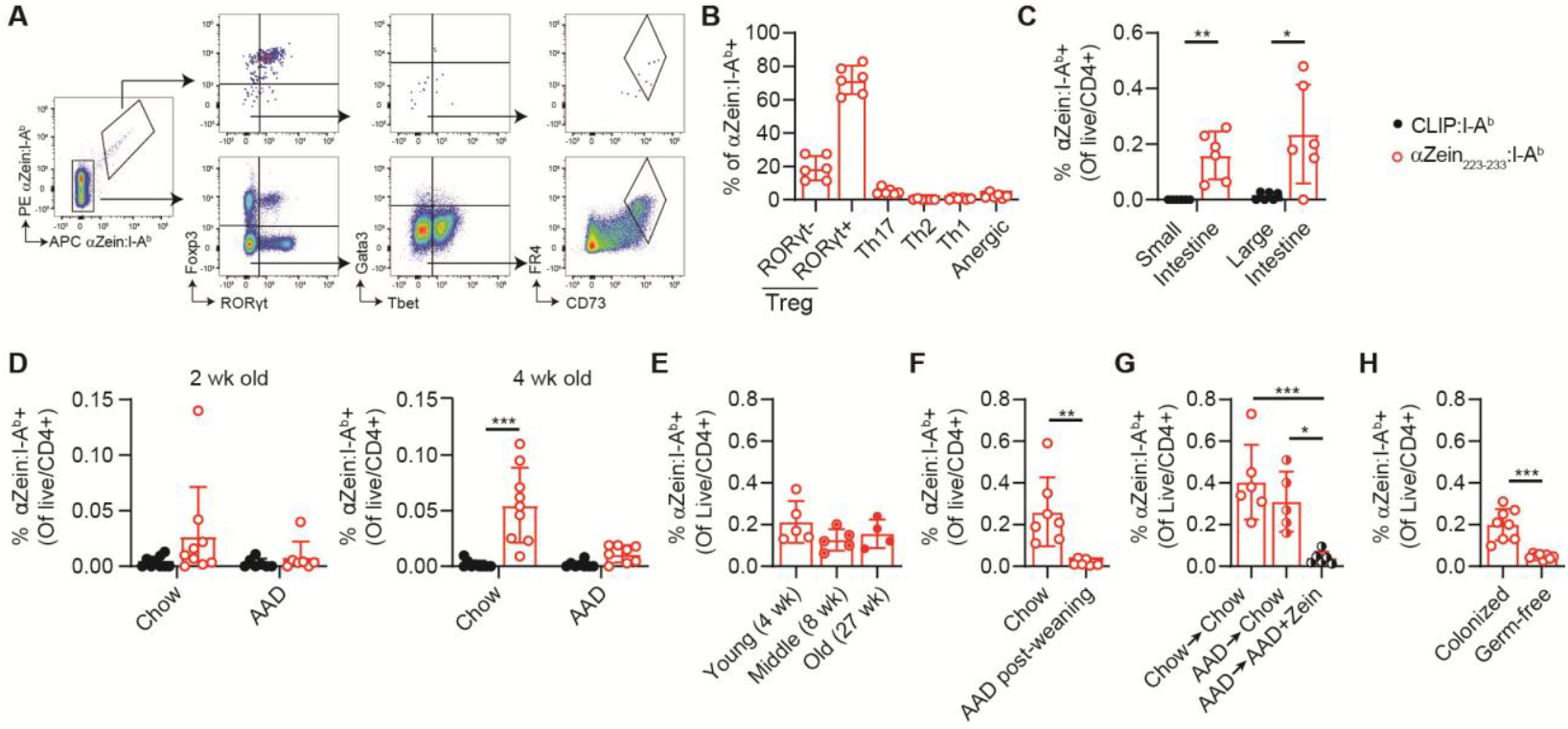
*In vivo* characterization and factors that influence the abundance of αZein_223-233_-responsive T cells. **(A-B)** αZein_223-233_-responsive T cells were profiled for T cell subtypes. RORγt^-^ Treg: Foxp3^+^RORγt^-^; RORγt^+^ Treg: Foxp3^+^RORγt^+^; Th17: Foxp3^-^RORγt^+^; Th2: Foxp3^-^RORγt^-^ Gata3^+^Tbet^-^; Th1: Foxp3^-^RORγt^-^Tbet^+^Gata3^-^; Anergic: Foxp3^-^RORγt^-^Tbet^-^Gata3^-^CD73^+^FR4^+^. N=6 mice/group. **(C)** Tetramer-positive cells in small and large intestine. N=6 mice/group. **(D)** Tetramer-positive cells in mice born onto chow or AAD diets profiled at 2 or 4 weeks of age. N=7-9 mice/group. **(E)** Tetramer-positive cells in mice at different ages. N=4-5 mice/group. **(F)** Tetramer-positive cells in mice swapped from chow onto an AAD diet at weaning or maintained on a chow diet, profiled at 12 weeks of age (9 weeks on AAD diet). N=7 mice/group. **(G)** Tetramer-positive cells in mice on chow diet or mice born on AAD diet then swapped onto chow or AAD+10% zein for 2 weeks, from 6-8 weeks of age. N=5-6 mice/group. **(H)** Tetramer-positive cells in colonized or germ-free adult mice. N=8 mice/group. All analyses are pre-gated on live CD4^+^ cells. (D) includes data from male and female mice, all other panels only use female mice. *P* values were calculated using a paired t-test (C), two-factor ANOVA (D) with a Sidak’s multiple comparisons test to follow-up on significant interaction terms, one-factor ANOVA (E,G) with a Tukey’s multiple comparisons test, or unpaired t-test (F, H). Unless otherwise indicated, all Tregs were analyzed from small intestine lamina propria of chow-fed mice. All mice from experiments in this figure were purchased from Jackson labs. Every dot represents an individual mouse. Error bars indicate mean ± SD. AAD, Amino acid defined. * p<0.05, ** p<0.01 and *** p<0.001.

While prior work has established the kinetics of pTreg cell induction post-birth, the development of food-responsive T cells is less well-understood, limiting our knowledge of the window during which tolerance develops. Food tolerance most likely arises either soon after birth, supported by the detection of food peptides in breast milk (*23*), or concomitant with weaning and the introduction of solid foods. To assess these possibilities, dams were randomized during pregnancy to a chow or an amino-acid-defined (AAD, antigen-free) diet. Their pups were fed the same diet until they were sacrificed at 2 or 4 weeks of age. Tetramer-positive T cells appeared at 4 wks of age only in the chow-fed mice (Fig. 3D, fig. S5A), mirroring an increase in total pTreg cells across weaning (fig. S5B-C). In other studies, pTreg cells were detected in the intestine 12 days after OVA exposure, suggesting that initial priming may have occurred around two weeks of age (*10*).

IgG antibodies against specific food components developed in parallel to food-responsive T cells, highlighting that food protein exposure activates both the cellular and humoral branches of the adaptive immune system (fig. S6A-B). These food-targeting antibodies constituted a significant portion of the total serum IgG, persisted throughout life contingent on continued chow exposure, and were also detectable in a panel of serum samples from healthy human donors (fig. S6C-E). While the role of IgG in oral tolerance, and relationship between Treg cells and antibodies is generally not well-understood, there is some evidence that IgG contributes to systemic food tolerance(*24, 25*). Together, these data show that the first introduction to food is a critical window for development of food-responsive peripheral Treg cells which appears to co-occur with an increase in IgG that recognize food components; both arms of immune recognition of food appear to reach a steady state that lasts through adulthood.

After weaning, the abundance and subtype distribution of αZein-specific T cells was stable across a range of ages (Fig. 3E, fig. S7A). As expected, this stability depended on continued chow exposure - mice swapped onto an AAD diet post-weaning experienced a sharp reduction in αZein-specific Treg cell abundance (Fig 3F). When mice born onto an AAD diet were introduced to chow post-weaning, the αZein-specific T cell response developed normally, though feeding a zein protein fraction without other chow components induced a much weaker response (Fig 3G, fig. S7B). This finding suggests a dependency on context of zein exposure (e.g., potential exposure to other food molecules and/or physical food matrix) for development of αZein-specific Treg cells, an aspect of food Treg cell development which is not shared with purified OVA (*10*). Further highlighting the importance of intestinal context, in germ-free mice the abundance of αZein-specific T cells was markedly reduced (Fig 3H), and phenotype was shifted toward fewer RORγt^+^ and more RORγt^-^ Treg cells (fig. S7C), revealing a potential interplay between immune development, gut microbiota, and food.

The identification of RORγt^+^Foxp3^+^ Treg cells specific for αZein suggests a possible parallel between food-responsive and bacterially induced T cells, since commensals such as *Helicobacter hepaticus* (*Hh*) induce the same ‘double-positive’ Treg cells (*26*). Induction of these *Hh*-responsive Treg cells depends on RORγt^+^ APCs (*27*). Similarly, we found MHCII^ΔRORγt^ mice, which lack MHCII expression on RORγt^+^ APCs, have fewer RORγt Treg cells and a corresponding increase in Th17 cells responsive to both αZein and gliadin (fig. S8). These results suggest similarities in requisite APCs between peripheral antigens of diverse origin (bacteria, food).

### Immunosuppressive factors are altered in food-responsive Treg cells

To search for molecular features that are characteristic of naturally-induced food-responsive T cells, we measured the transcriptional profile of αZein-specific Treg cells. In one experiment, we used single cell RNA-sequencing to profile αZein-specific Treg cells, Helicobacter-responsive Treg cells, and bulk controls (all Treg cells regardless of antigen-specificity) (fig. S9-11). In a parallel experiment, bulk RNA-sequencing was used to compare αZein-specific Treg cells to adoptively transferred OVA-specific Treg cells (following exposure to OVA in drinking water), and bulk Treg cells (all CD4^+^CD25^+^ cells of unknown antigen specificity) from chow-fed, AAD-fed, or germ-free (GF) mice (fig. S12).

Several genes associated with immune suppression (e.g., *Gzmb* and *Lag3*) emerged from the sequencing data as associated with food-responsiveness (Fig. 4A, data file S4). Follow-up flow cytometry assays revealed distinct profiles of immune suppressive markers (e.g., Lag3, Gzmb, Tim3, CTLA4) in αZein-specific Treg cells, other food-antigen specific Treg cells, and across bulk Treg cell populations from mice with different environmental antigen exposures (AAD diet, germ-free) (Fig 4B-D, fig. S13-15). Recently, some of the same immune suppressive genes were also identified as upregulated in intestinal epithelium resident regulatory T cells from chow compared to AAD diet fed mice, suggesting a conserved effect in multiple intestine layers (*7*). Our tetramer data corroborates this finding and specifically implicates food-responsive T cells as a source of these differences. We considered activation status as a possible explanation for the observed differences, either reflecting underlying Treg cell biology or as an artifact of tetramer staining. Prolonged tetramer staining can lead to T cell activation, thus altering phenotype (*28*); however, experiments with OVA suggested that our tetramer staining protocol did not alter immune suppressive molecule profile (fig. S16). Further, our single cell RNA-sequencing data did not show enrichment of canonical Treg cell activation markers in αZein-specific Treg cells compared to bulk controls, and across all Treg cells there was no apparent relationship between multiple immune suppressive and activation associated transcripts (fig S17A-B) (*29*–*31*). While activation status at the time of cell isolation is unlikely to explain differences in transcript profile, intentional ex vivo activation did exacerbate differences in immune suppressive programs between αZein-specific and bulk Treg cells. Lag3 expression was induced in intestinal αZein-specific Treg cells by chemical (PMA/ionomycin) or peptide stimulation and diminished following a short-term Zein washout (3 days on AAD diet) (Fig 4E-F). These data reveal that the transcriptional profile of αZein-specific Treg cells is dynamic and influenced by a combination of factors, including recency of antigen exposure and activation by external elicitors.

**Fig. 4:**
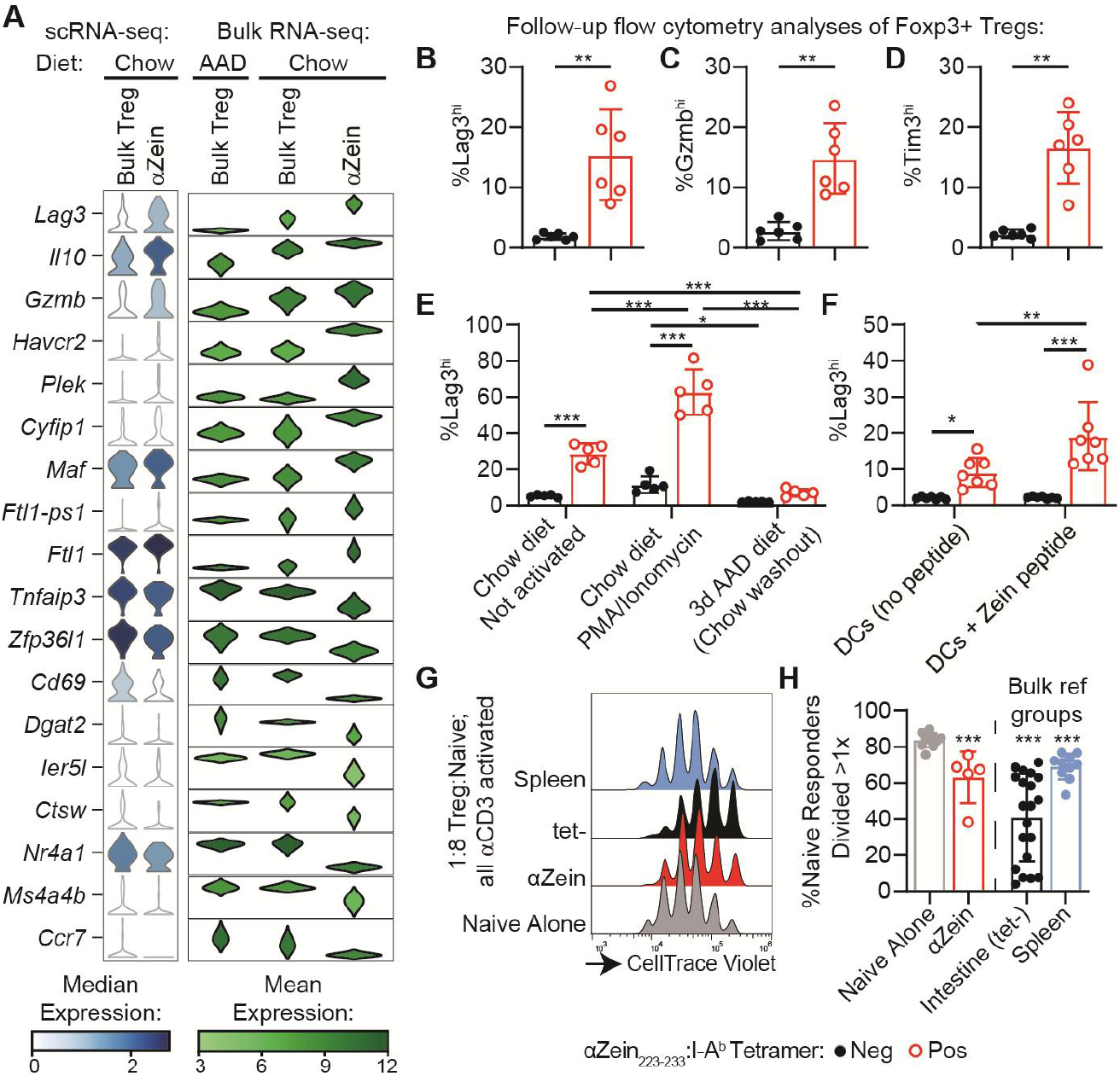
Food-responsive Treg cells display a distinct transcriptional signature of immune suppressive markers. **(A)** Violin plots showing a selection of differential genes of interest shared in both the single cell and bulk sequencing datasets. Single cell data compare bulk Tregs from chow fed control mice to antigen-specific αZein_223-233_ Tregs and the bulk sequencing data compare bulk Tregs from chow fed control mice, AAD fed mice, or antigen-specific αZein_223-233_ Tregs. **(B-D)** Lag3, Gzmb, and Tim3 levels between αZein_223-233_ binding and tetramer non-binding Tregs. N=6 mice/group. **(E)** Lag3 levels between αZein tetramer-positive and -negative populations, under control conditions, activation by a 4 hour PMA/ionomycin stimulation, or 3 days on AAD diet compared to control chow fed mice. N=5 mice/group. **(F)** Lamina propria single cell suspension co-cultured for 4 hours with dendritic cells (no added peptide) or dendritic cells pulsed with zein peptide (FYQQPIIGGAL). N=7 mice/group. **(G-H)** Percent of naïve T cells divided after incubation with antigen presenting cells, a Treg population (either αZein-specific Tregs, tetramer-negative Tregs, or splenic Tregs), and αCD3 antibodies. N=10 naïve controls, 5 Zein Tregs, 20 intestinal tet-Tregs, and 10 spleen Treg samples. Statistical analysis was used to compare each group against the Navie alone control. (B-F) are gated on live CD4^+^Foxp3^+^ and αZein:I-Ab-PE^+^αZein:I-Ab-APC^+^ (tetramer-positive) or αZein:I-Ab-PE^-^αZein:I-Ab-APC^-^ (tetramer-negative). (G-H) are gated on live CD4^+^CD45.1^+^CTV^-^. *P* values are calculated using paired t-test (B-D), or two-factor ANOVA (E-F) with a Tukey’s post-hoc test or Uncorrected Fisher’s LSD test, or unpaired t-test (H). Every dot represents an individual mouse. Error bars indicate mean ± SD. AAD is amino acid defined diet. * p<0.05, ** p<0.01 and *** p<0.001.

To further assess cell state, we compared our single cell sequencing data against published Treg cell gene expression profiles. Compared to bulk intestinal Treg cells, αZein-specific Treg cells aligned more closely to tissue-adapted ‘effector Tregs’ or ‘IL10^stable^ Treg cells’, suggesting that habitual zein exposure induces a stable tissue-resident Treg cell population (*32, 33*) (fig. S17C-E). We tried to leverage the observed single cell gene expression profile to identify additional food-responsive TCRs. This effort revealed an additional Gliadin_273-285_ responder, highlighting the immunodominance of this epitope and some capacity for gene expression profile to reveal antigen specificity. However, this approach was not a systematic improvement over other mapping strategies (Table S1, fig. S18).

To more directly assess the suppressive capacity of food responsive Treg cells, we sought to measure αZein-specific Treg cell-mediated suppression of naïve T cell expansion following an ex vivo αCD3 stimulation. Isolated αZein-specific Treg cells effectively suppressed division of naïve T cells (Fig 4G-H). We observed that the suppressive capacity of the bulk Treg cell pool was highly variable and also changed depending on diet and anatomical site, likely reflecting the distinct arsenal of immune suppressive factors expressed by each population (fig. S19-20). For example, while chow Treg cells from the lamina propria were Lag3^hi^ and Gzmb^hi^, consistent with our RNA-sequencing data, these cells simultaneously expressed low levels of other immune suppressive makers such as Klrg1 and PD-1. Taken together, αZein-specific Treg cells adopt a phenotype reflective of a mature tissue-resident cell state and enriched in select immune suppressive factors. These data provide direct evidence that Treg cells responding to an immunodominant food epitope engage in canonical suppression programs (*34*–*36*), and that a single Treg cell can concurrently use multiple suppressive strategies. Furthermore, these suppressive programs appear to be upregulated by recognition of the cognate food epitope.

### Oral consumption elicits systemic zein tolerance

We next evaluated contexts in which αZein-specific Treg cells are immunomodulatory in vivo. First, we measured antibody levels following an inflammatory intraperitoneal sensitization, a model which robustly reveals tolerance in mice previously exposed to oral OVA (fig. S21A-B). Surprisingly, no anti-zein antibodies were observed after intraperitoneal zein sensitization in mice consuming either a defined corn-containing (AAD + corn/soy/wheat/oat) or corn-free (AAD + soy/wheat/oat) diet (fig. S21C-D). When mice on the same diets were exposed to intraperitoneal OVA, OVA antibodies were robustly detected (fig. S21E). To explore this further, we injected chow or AAD fed mice with whole corn extract (fig. S22A). In this model, both corn- and zein-targeted antibodies were higher in AAD mice compared to chow-fed mice, consistent with a protective effect of chow (fig. S22B-C). Moreover, MHCII ^ΔRORγt^ mice, that had fewer RORγt^+^ Treg cells responsive to αZein, also showed higher levels of corn- and zein-targeted antibodies compared to wildtype controls despite all mice consuming chow diet (fig. S22D-F). Collectively, these data reveal that in conditions where αZein-specific Treg cell development is impaired, either through diet or genetic knockout, oral tolerance to systemic challenge is also less effective. To further investigate oral tolerance to a complex lysate, a similar model was used to probe oral tolerance toward sesame (Fig. S22G). Given that we observed cross-reactivity between the TCR we identified as reactive to Glycinin G1 with sesame lysate, we included a soy containing diet as a control group. Indeed, sesame consumption induced oral tolerance to sesame and soy consumption provided protection compared to an AAD diet, revealing potential protective effects of antigen cross-reactivity (Fig. S22H). However, we cannot rule out a role for diet complexity or a diet containing any protein in explaining protective effects of soy.

As an orthogonal approach to measuring zein allergy, we employed a cholera toxin driven oral sensitization model. This model robustly revealed OVA allergy, but provided no evidence of zein or gliadin allergy in mice born on an AAD diet (fig. S23A-D). To better understand the relationship between antigen and allergy development in this model, we further tested casein and lactoglobulin, two milk proteins with different biochemical properties. While some of the lactoglobulin sensitized mice experienced anaphylactic allergy, none of the casein sensitized mice displayed symptoms (Fig S23E-H). These data suggest that inflammatory responses to proteins (including zein) depend on the context and perhaps also the identity of the protein. However, these data also emphasize a protective effect of oral corn exposure to subsequent immune responses to corn, consistent with an immune suppressive nature of αZein-specific Treg cells.

To more directly probe the function of T cell-mediated zein tolerance, we employed a CFA-driven inflammation model (*37*). Injection of zein epitope emulsified in CFA induced expansion of αZein-specific T cells in the inguinal lymph node (ILN) (Fig. 5A). To determine the role of oral tolerance in modulating this response, we compare mice born onto chow diets with existing zein tolerance to mice born onto AAD diets for which this was the first zein exposure (Fig 5B). Consuming chow diet reduced the frequency of the CFA-induced Zein-responsive T cells in the draining lymph and skewed the T cell response to include more Treg cells and fewer anergic cells (Fig. 5C-D), consistent with a tolerogenic immune suppressive response. This effect was dependent on dietary exposure to the specific epitope, as evidenced by the observation that the T cell response to a non-dietary epitope (OVA) epitope was the same on chow or AAD diet (Fig 5E-F). Correspondingly, when isolated lymph node cells were stimulated with zein peptide *ex. vivo*, IL-2 production, a marker of T cell activation, was constrained in samples from the chow fed mice (fig. S24A). Dietary background did not affect T cell activation for an unrelated antigen (2W1S, fig. S24B). To directly assess the function of the Zein-responsive Treg cells from chow fed mice, we tested their immune suppressive capacity against naïve T cells. Indeed, CFA-expanded αZein-specific Treg cells isolated from the inflammatory lymph node setting displayed robust immune suppression (Fig 5G-H). Further, these lymph-derived αZein-specific Treg cells from chow fed mice displayed a Lag3^hi^ phenotype comparable to that observed in intestinal αZein-specific Treg cells (fig. S24C). Blocking Lag3 alleviated some of the αZein-specific Treg cell-mediated suppression, suggesting a causal role for Lag3 in αZein-specific Treg cell-mediated suppression (fig. S24D-F).

**Figure 5:**
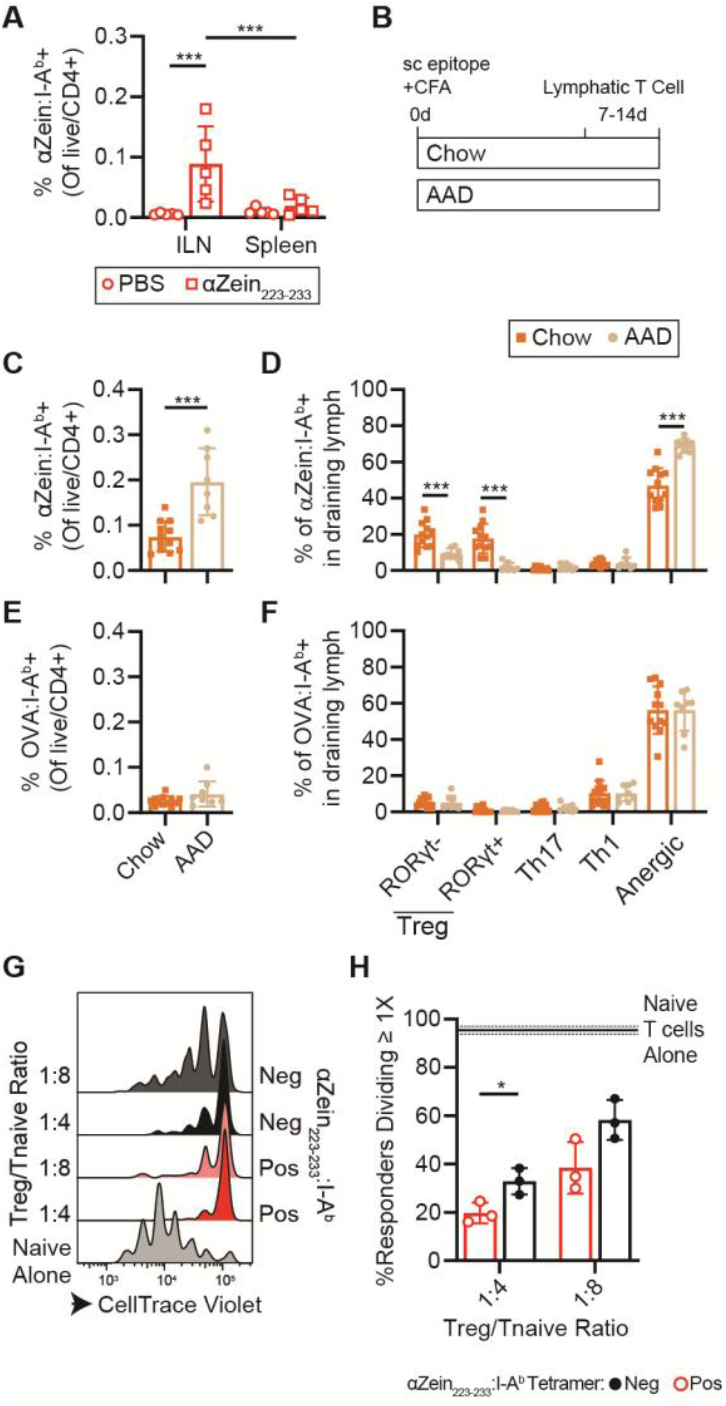
αZein-specific T cell profile and suppressive capacity after CFA simulation. **(A)** Tetramer-positive cells in the inguinal lymph node (ILN) and spleen following an intraperitoneal injection with CFA or zein epitope emulsified in CFA. N=5 mice/group. **(B)** Mice were born onto chow or AAD diets. In adulthood (6-10 wks) mice were given a subcutaneous injection of zein epitope emulsified in CFA on day 0. T cells were isolated from the draining lymph node between 8-14d post-injection. **(C-F)** Abundance and distribution of cell types responsive to αZein or OVA in draining lymph nodes following epitope +CFA injection. N=8-12 mice/group. Only significant pairwise comparisons within a cell type are shown. **(G-H)** Percent of naïve T cells divided after incubation with antigen presenting cells, a Treg population (αZein-specific Tregs or tetramer-negative Tregs isolated from the inguinal lymph node) and αCD3 antibodies. N=3 mice/group. (C) and (E) are gated on live CD4^+^TCRβ^+^, (D) is gated on live CD4^+^TCRβ^+^αZein:I-Ab-PE^+^αZein:I-Ab-APC^+^, and (F) is gated on live CD4^+^TCRβ^+^OVA:I-Ab-PE^+^OVA:I-Ab-APC^+^. *P* values were calculated using a two factor repeated measures ANOVA with an Uncorrected Fisher’s LSD test (A), an unpaired t-test (C,E,H), two-factor repeated-measures ANOVA with a Sidak’s multiple comparisons test (D), or two-factor repeated-measures ANOVA (F). Every dot represents an individual mouse. Error bars indicate mean ± SD. * p<0.05, ** p<0.01 and *** p<0.001.

As a final measure of Treg cell function, we adoptively transferred αZein-specific Treg cells into a naïve recipient mouse and measured the sufficiency of these Treg cells to mediate immune tolerance. To achieve necessary numbers of αZein-specific Treg cells and total cells for transfer, we used CFA-expanded (with or without zein epitope) total CD4^+^CD25^+^ Treg cells from chow-fed mice as the donor cell population (fig S25A-B). The activity of these Treg cells was then tested in the recipient mice using a subcutaneous epitope + CFA challenge model, with T cell abundance and phenotype as the readout. This model was adapted from a similar assay previously used to show tolerance toward self-antigens, and enabled discrimination of adoptively transferred and host T cells (*38*). Transfer of αZein-specific Treg cells suppressed the generation of αZein-specific Th1 cells in recipients (fig. S25C-F). The frequency of newly induced Zein-specific T cells and IL-2 secretion in response to *ex vivo* restimulation was unaffected by the presence of adoptively transferred αZein-specific Treg cells, which could suggest specific suppression of Th1-driven inflammation in this model (fig. S25G-H). Overall, these data reveal that zein tolerance achieved through chow consumption facilitates Treg cell-mediated suppression of a future inflammatory challenge toward zein.

## DISCUSSION

Oral tolerance is a remarkable process of suppressing immune responses toward dietary proteins. Our ability to characterize the development of oral tolerance and mechanisms of Treg cell-mediated immune suppression have been limited because the antigens that are natural Treg cell ligands were unknown. Here, we identified food epitopes that are recognized by naturally-induced Treg cells after oral introduction of antigens. The antigens identified – αZein, glycinin, and gliadin – are all seed storage proteins, suggesting that this class of proteins is a common source of tolerogenic epitopes. We find that αZein-specific Treg cell emergence depends on several factors including developmental state and intestinal context (microbiome, food matrix, sampling mechanisms). Further, we demonstrate that food-responsive Treg cells are characterized by a distinct transcriptional profile, including upregulation of a subset of immune suppressive molecules. These immune suppressive agents were previously reported as differential across bulk Treg cell populations from chow and AAD fed intestinal epithelial or lamina propria cells, but not differential between a gut microbe-specific Treg cell population and bulk controls, confirming a specific association with food-reactive Treg cells (*3, 7, 39*). Lastly, we demonstrated the functional immune suppressive capacity of αZein-specific Treg cells and their role in oral tolerance. Notably, oral exposure to corn under conditions that elicit αZein-specific Treg cells protects against an inflammatory response to corn driven by IP immunization. Adoptive transfer experiments with αZein-specific Treg cells revealed the specific role of these cells in promoting oral tolerance. We anticipate our findings regarding oral tolerance toward zein, as an abundant and stable dietary protein, will directly translate to other seed storage proteins, including those that are prevalent in food allergy.

Our characterization of zein tolerance, combined with other model studies of oral tolerance, begins to highlight potential contributions of the specific antigen to observed immune outcomes. High dose OVA in drinking water coupled to adoptively transferred OTII cells is a robust and widely used oral tolerance model. In contrast, dietary zein alone only weakly induced Treg cells compared to zein incorporated in chow. Intraperitoneal OVA, not but zein, strongly induced antibodies. These data suggest a possible role for food matrix in promoting immune recognition of zein through both the oral and intraperitoneal routes. T cell phenotype is also antigen- and context-dependent. For example, while chow consumption induced a predominant αZein-specific Treg population, after 1 week on a gliadin containing diet, many gliadin-responsive T cells adopted an anergic phenotype (*6*). In other studies, after short term OVA exposure, only half of adoptively transferred OVA-specific T cells become Foxp3^+^ Tregs (*3, 10*). Determining whether these differences reflect antigen-intrinsic properties or experimental parameters (dose, duration, age at initial exposure, etc.) is essential to understand the range of tolerance phenotypes. Treg populations induced by short-term feeding could be functionally distinct from long-term gut-resident Treg cells, and the functional consequence of phenotype for durable tolerance is unknown. Thus, identifying and characterizing chow-responsive T cells, is essential to understand the nature of long-standing oral tolerance.

It is notable that the food Treg cell epitopes we have found from two different common dietary grains - including the immunodominant tolerance antigen from corn - derive from highly abundant, water-insoluble proteins. Furthermore, corn is widely tolerated in human populations and does not commonly result in food allergy. Our data suggests that this lack of a negative immune response is not due to limited access of corn proteins to intestinal immune cells. In contrast, our measurement of αZein-specific Treg cells indicates that corn proteins are effectively sampled and presented by gut-resident APCs. In contrast to the proteins we identified as sources of Treg cell antigens, proteins that are known to be recognized by Th2 cells in food allergy are largely water soluble. Together, these observations further underscore that physical properties coupled with relative abundance of key proteins in food might directly impact whether a given food is likely to be widely tolerogenic versus susceptible to immune sensitization. Further experiments are needed to determine whether solubility or other biochemical factors are key determinants of pathogenic immunogenicity. This unexpected outcome of our work is worth future investigation as it could help explain why some foods are disproportionately associated with increased incidence of food allergies. Identifying tolerance epitopes has direct implications for understanding allergies. For example, Glycinin G1 is a known allergen (Gly m 6). However, the epitope we identified does not correspond to a known allergen epitope (from the immune epitope database (IEDB), (*17, 40*–*46*)), with the caveat that IEDB consists primarily of human epitopes. In addition, the glycinin-responsive TCR is not activated by known Gly m 6 cross-reactive lysates (e.g., peanut lysate/allergen Ara h 3 (*47, 48*)). In the future, tracking T cell responses to this epitope under inflammatory or homeostatic conditions may reveal vulnerabilities that initiate allergy development. Further, understanding how proteins like αZein, that are not common allergens, drive strong Treg responses, could provide a blueprint for programming antigen-specific Treg responses. Cereal grains comprise over half of the world’s daily caloric intake, and we anticipate that the proteins identified here will also be common targets of Treg cells in humans (*49*). Indeed, our analysis of human serum samples begins to provide evidence for immune recognition of multiple dietary grains, including corn.

Diet is our most intimate interaction with our environment. Correctly recognizing foods as safe creates an anti-inflammatory environment to support nutrient acquisition and prevent allergy. This research advances our understanding of the major dietary antigens recognized by intestinal Tregs and demonstrates their function in oral tolerance toward prolamin antigens. Tapping mechanisms of oral tolerance as naturally occurring programming with molecular specificity could enable the use of synthetic approaches for redirecting of allergic and autoimmune states.

## MATERIALS AND METHODS

### Study design

This study was designed to identify antigens from mouse diet that are recognized by regulatory T (Treg) cells following homeostatic chow consumption. First, we generated hybridoma cell lines bearing TCRs identified on Treg cells from intestinal single-cell RNA-sequencing datasets. These hybridomas were screened using an *in vitro* mixed lymphocyte assay. For receptors that were reactive to a food component, we mapped the specific antigen epitope and determined cross-reactivity toward other plant extracts. One epitope from the maize protein alpha-zein emerged as immunodominant and was prioritized for further study. An MHCII tetramer loaded with this epitope was used to characterize the abundance, location, development, and persistence of food-responsive Treg cells. Zein-reactive Treg cells were further characterized using both bulk and single-cell RNA-sequencing. Lastly, we used numerous in vitro and in vivo models to assess the capacity of αZein-specific Treg cells to perform immune suppression and mediate oral tolerance. Sample sizes were determined based on prior experience. Mice were randomized to studies and no mice were excluded. Blinding was not performed.

### Tissue samples from mouse and human

All studies were conducted under administrative panel for laboratory animal care approved protocols at Stanford (Protocol 33997) or institutional animal care and usage committee protocols through the New York University School of Medicine (NYU) and adhered to ethical standards for the treatment of research mice. Animals were housed in a conventional facility with 12h light-12h dark cycles. When indicated, mice were fed amino acid defined AIN-93G diet (Dyets Inc, Item 510017). Experiments at Stanford were performed with C57BL/6 animals from Jax (Strains: 002014 or 000664) For the development study, timed pregnant C57BL/6 mice were purchased from Jax. Mice were randomly maintained on chow diet or switched onto the amino acid defined (AAD) diet upon arrival and gave birth within one week. For all other studies of mice born on AAD diets, C57BL/6 mice were bred in-house on AAD diet starting when breeders were set up. When indicated, custom diets were made by mixing L-AA defined AIN93G diet with other human-grade food ingredients at the indicated percentages. Germ-free mice were maintained in aseptic incubators.

Ptprc^a^Pepc^b^/BoyJ mice were purchased from Jax and given 1 week to acclimate to the animal care facility. On the first day of the study, 1x10^6^ OT-II cells were adoptively transferred from Rag2/OT-II mice (Strain 11490, Taconic) by retroorbital injection. Mice then received 10 mg/ml OVA (Sigma) in drinking water for 7 days. At NYU Grossman School of Medicine, all transgenic mice were bred and maintained in the Alexandria Center for Life Sciences – West Tower vivarium in specific pathogen-free conditions. C57BL/6 mice (Jax#000664), I-AB^f/f^ (B6.129X1-H2-Ab1tm1Koni/J Jax 013181) mice were purchased from Jackson Laboratories. RORγt-*cre* and Hh-72tg were generated by members of the Littman laboratory and have previously been described (*26, 50*). Female and male mice were used equally in this experiment. Mice from 6-12 weeks of age were used. All mice were housed with a 6AM-6PM light on-off cycle with an ambient temperature of 18-24 degrees Celsius and humidity maintained between 30-70%. Human serum samples were obtained from the Stanford Blood Center from anonymous donors.

### Intestinal T cell isolation

Small intestine was removed from the animal and adipose tissue and Peyer’s patches were carefully removed. A sagittal cut was made through the lumen, opening the intestine into a flat layer. Intestinal samples were incubated at 37°C in a solution of Dulbecco Modified Eagle Medium [DMEM (VWR)]+ 5% fetal bovine serum [(FBS), Thermo Fisher] with 5 mM EDTA (Sigma Aldrich) and 1 mM DTT (Fisher) for 40 minutes with shaking. Next, samples were placed in conical tube with warm DMEM and shaken vigorously for ∼20 seconds. Samples were collected on a strainer and the shaking step was repeated once more. Samples were then processed on a gentleMACS (Miltenyi) using the LPDK-1 program and enzyme solutions from the lamina propria dissociation kit (#130-097-410, Miltenyi). When samples were finished on the MACS, an equal volume of 80% Percoll PLUS density gradient media (Cytiva) was added, diluting the sample to 40% percoll. The sample was moved to a new conical tube, and 80% percoll was added to the bottom using a transfer pipette, resulting in two layers. The samples were centrifuged at 600 x g for 10 minutes at room temperature, and cells were recovered from the interface of the 40 and 80% percoll layers. Cells were washed in 10 ml of media and then passed through a filter prior to preparing for flow cytometry analysis.

### Flow cytometry

All tetramers were provided by the NIH Tetramer Core Facility with both PE and APC fluorophores. The following tetramers were used: αZein (FYQQPIIGGAL), CLIP (PVSKMRMATPLLMQA), Gliadin (NVYIPPYCTIAP), and OVA (AAHAEINEA and HAAHAEINEA). The two OVA tetramers were used as a pool. Tetramer staining was performed for 1 hour at 37°C in Roswell Park Memorial Institute ([RPMI], Thermo Fisher, 61870127) media with 10% FBS or for 1 hour at room temperature in MACS buffer with 5% FBS at a 1:100 dilution. To enhance detection of antigen-specific T cells in the CFA-peptide experiment, cells were treated with a protein-kinase inhibitor—dasatinib (MedChemExpress, HY-10181)—at a final concentration of 50 nM for 30 min prior to and throughout tetramer staining at 37°C. Cells were stained with viability dye, and in some experiments cell surface markers, in MACS buffer (Miltenyi Biotec) with 5% FBS, then fixed and permeabilized using the eBioscience Foxp3/Transcription factor staining buffer set (ThermoFisher, 00-5523-00). Cells were fixed for either 1 hour at room temperature or overnight at 4°C. Intracellular targets, and in some experiments intracellular and cell surface targets were stained in permeabilization buffer. All antibodies used in are indicated in Table S2. At Stanford, cells were analyzed using an LSRII or Symphony (BD Bioscience) analyzer and sorted using a FACSAria II at the Stanford Shared FACS Facility. Flow cytometry experiments at NYU were performed on an Aurora (Cytek) analyzer, while cells were sorted using a FACsAria (BD Bioscience). All flow cytometry data was analyzed using FloJo software.

### Hybridoma generation

Hybridomas were generated as previously described (*13*). Briefly, pMSCV-mCD4-PIG TCR-OTII backbone was used in combination with TCRα/TCRβ sequences of interest (data file S5), which were separated by a P2A peptide and synthesized commercially (Twist Bioscience) or cloned in lab. αZ_TCR_1, αZ _TCR_2, αZ _TCR_3, and Gly_TCR_1 had been generated and found to be food responsive previously (*13*). All other hybridomas were generated in this study. Lipofectamine 3000 reagent (L3000001, Thermo Fisher) was used to generate virus particles containing the TCR vector in Platinum-E cells (RV-101, Cell Biolabs). The virus was then used to transduce NFAT-GFP hybridoma cells. The presence of a functional TCR in all hybridomas was confirmed by measuring IL-2 secretion after stimulation with an anti-CD3 antibody. Some hybridomas were selected based on clonal relationship identified using GLIPH2 (*51*).

### Treg suppression assay

CD45.2+CD4+GFP+αZein_223-233_:I-Ab+ Tregs and CD45.2+CD4+GFP+αZein_223-233_:I-Ab-Tregs were sorted from the small intestine lamina propria of Foxp3-GFP reporter mice (B6.Cg-Foxp3tm2Tch/J, Jax Strain, 006772). CD45.2+CD4+GFP+ Tregs were also sorted from spleens. For some experiments, Tregs were instead isolated from draining lymph nodes 11d following CFA-peptide sensitization, following the methods described below. Naïve CD45.1+ T cells were isolated from spleen by pre-enrichment using a Naïve CD4+ T cell isolation kit (Miltenyi), stained with CellTrace Violet dye (CTV, Thermo Fisher), and subsequently purified by flow cytometry (live/CD4+/CD62L+/CD44-). Tregs were cultured at a ratio ranging from 1:32 to 1:2 (determined based on number of Tregs sorted) with naïve T cells in the presence of 1ug/mL anti-CD3 (Biolegend) and dendritic cells (DCs, isolated as described above) at a 1:1 DC-to-T cell ratio in a V-bottom plate. Naïve T cells cultured alone with anti-CD3 and DCs were used as a control. In some experiments, cells were treated with 30 ug/ml anti-Lag3 (BioXCell, BE0174) or 30 ug/ml anti-Isotype control (BioXCell, BE0088). Cell proliferation was measured after 3 days of co-culture by measuring CTV dilution in the CD45.1+ T cell population. During the assay, cells were cultured in RPMI 1640 + 10% FBS + 1% penicillin-streptomycin + 1X Sodium Pyruvate (Gibco) + 1X MEM Non-essential amino acids (Gibco) + 50 µM BME (VWR).

### Intraperitoneal allergy models

For measuring OVA tolerance, C57Bl/6 mice were given water (not tolerized) or water supplemented with 10 mg/mL OVA (Sigma) (tolerized) *ad libitum* for 1 week. Two days after ceasing supplemented water feeding, both groups of mice were intraperitoneally injected with 100 uL of a solution containing 50 ug OVA precipitated with 1 mg Imject Alum (Thermo Fisher) in PBS. Mice were given a booster dose in a similar fashion 2 weeks after the initial injection with 100 uL of a solution containing 50 ug OVA in PBS. Serum was collected one week after the second injection and assayed for anti-OVA IgG1 at a 1:6,000 dilution using a commercially available ELISA kit (Cayman).

For zein tolerance, in the first experiment, mice had three dietary backgrounds: born and maintained on chow; born on AAD and randomized at 4 wks of age to start a defined corn-containing diet (AAD diet + 10% each of corn, soy, wheat, oat); born on AAD and randomized at 4 wks of age to start a defined corn-free diet (AAD diet + 10% each of soy, wheat, oat). At 6 wks mice began the inflammatory injection series which consisted of an initial dose of 500 ug protein + 1 mg of Imject Alum (Thermo Fisher) on day 0, a booster dose of 50 ug protein on day 14, and serum collection post-sacrifice on day 21.

In the parallel experiment to measure zein tolerance, mice born onto chow or AAD diets were sensitized with IP corn. At 6 wks of age mice began the inflammatory exposure series consisting of a 500 ug dose of corn +1 mg of Imject Alum (Thermo Fisher) on day 0, booster with 50 ug corn on day 14, and serum collection post-sacrifice on day 21. The same experimental series was repeated with MHCII ^ΔRORγt^ mice and control mice fed chow diets.

### Oral allergy model

To induce allergy, mice were sensitized to zein (Santa Cruz), gliadin (Sigma), OVA (Sigma), casein (Santa Cruz) or β-lactoglobulin (Sigma) alongside 10 ug of cholera toxin (Millipore) in 200 ug 5% Sodium bicarbonate 2 times (days 0 and 7). On day 14, mice were challenged with an intraperitoneal injection of 2 mg of cognate allergen. Body temperature was measured every 5 minutes for 1 hour using the BMDS transponder system.

### Peptide administration in CFA

In the initial experiment mice were injected intraperitoneally with 100 ug of zein epitope emulsified in CFA. The spleen and inguinal lymph node were harvested after 7 days. In subsequent experiments, mice were subcutaneously injected in the right flank or tail base with 100 uL of CFA (Invivogen) emulsion containing 10 ug of each epitope of interest. Peptides (2W1S: EAWGALANWAVDSA; OVA: ISQAVHAAHAEINEAGR; Zein: FYQQPIIGGAL) were purchased from Genscript. Draining inguinal and axillary lymph nodes were harvested 14 days post-injection and processed into single-cell suspensions which were then stained with relevant tetramers and antibodies specific for phenotypic markers as described previously. Draining inguinal and axillary lymph nodes were harvested 14 days post-injection and processed into single-cell suspensions which were then stained with relevant tetramers and antibodies specific for phenotypic markers as described previously. In experiments to probe the suppressive capacity of Zein Tregs, only the zein epitope was injected. As an orthogonal readout of activation, mice were injected with 10 ug of Zein or 2W1S epitope. On day 8, single cell suspensions were isolated from the inguinal lymph node and restimulated ex vivo with 10 ug/ml of cognate peptide. After 24 hours, media IL-2 levels were measured using an ELISA assay as described above.

### Statistics

Statistical analysis was performed in GraphPad Prism 9. Comparison of two groups was performed with a t-test and comparison of 3 or more groups with a 1-factor analysis of variance (ANOVA). Data with two variables was analyzed using a 2-factor analysis of variance (ANOVA). When data were collected from paired samples, for example tetramer-positive and negative data from the same mouse, a paired t-test was performed. Tukey’s multiple comparisons test, Dunnett’s multiple comparison test, Sidak’s multiple comparisons test, or the Uncorrected Fisher’s LSD test were used for post-hoc analysis of ANOVA data. P < 0.05 was considered statistically significant.

## Supporting information

datafile_S1

datafile_S5

datafile_S6

datafile_S4

datafile_S2

datafile_S3

## Supplementary Materials

**The PDF files includes:**

Methods

Table S1-S2

Figs. S1-S25

## Other Supplementary Material for this manuscript includes the following

Data files S1-S5

MDAR Reproducibility Checklist

## ACKNOWLEDGEMENTS

We thank Kevin O’Connor and Mydia Phan for assistance with tissue collection for the antigen presenting cell elucidation and single cell sequencing experiments; Steven Higginbottom for providing germ-free mice; Henry Le for helpful discussions and manuscript feedback; and Diego Wengier, Alexa Weingarden, and Djenet Bousbaine for helpful discussions. We thank the NIH Tetramer Core Facility (contract number 75N93020D00005) for providing the MHCII tetramers.

## Author contributions

JEB, KN, MAF, ESS conceptualized the project; JEB, RK, KN, MAF, ESS, DRL designed experiments; JEB, RK, KN, EAS, FMC, RU, GRM carried out the experiments; JEB wrote the paper with edits from ESS, RK, EAS, KN, FMC, RU, DRL, MAF.

## Data, code, and materials availability

Sequencing data generated in this study is deposited in EMBL-EBI ArrayExpress under accession number E-MTAB-16143 (bulk RNA-sequencing) and E-MTAB-16270 (scRNA-sequencing). Tabulated data underlying all the figures are provided in data file S6. All other data needed to support the conclusions of the paper are present in the paper or the Supplementary Materials. Code is available at Zenodo (*52*). Tetramer reagents unique to this research were provided by the NIH Tetramer Core Facility (NIH Contract 75N93020D00005 and RRID:SCR_026557) and are available from them upon request. Hybridoma cell lines were generated by the authors and are available from J.B. under an MTA from the Salk Institute. RORγt-*cre* and Hh-72tg mice are available from Jax (Stock 032538 and 022791). All other materials are commercially available as described in the Materials and Methods section.

