## Supplemental Information for "Identification and characterization of dietary antigens in oral tolerance"

**The PDF file includes:**

Materials and Methods
Figs. S1 to S25
Tables S1 to S2

### Supplementary Materials and Methods

Lymph node isolation:

Mesenteric lymph nodes were removed from mice, gently mashed with a syringe plunger through a 100 um strainer, and rinsed with 10 mL of RPMI 1640+10% FBS. The resulting suspension was centrifuged at 500xg for 5 minutes, resuspended in RPMI 1640+10%FBS, and filtered through a 35 um FACS tube strainer to remove large aggregates.

Preparation of antigen formulations:

The mouse chow (5K67 diet) fed to the discovery cohort contains 7 food ingredients: wheat, corn, oat, fish meal, soy, alfalfa, yeast. Ingredients were purchased from Amazon and ground in PBS using a ball mill homogenizer (Retsch) or mortar and pestle. In follow-up studies, seeds from plants of interest were purchased from commercial sources (Amazon, plant nurseries, online retailers) and similarly ground in PBS. Synthetic peptides were purchased from GenScript.

Dendritic cell isolation:

Mice were injected with 5 x $10^6$ B16-FLT3L cells by intraperitoneal injection to induce spleen hypertrophy and dendritic cell population expansion, as previously described (13). After sacrifice, spleens were excised, and a single cell suspension was obtained using a spleen dissociation kit (Miltenyi Biotec, #130-095-926) the gentleMACS dissociator (Miltenyi Biotec). After the dissociation, red blood cells were lysed (RBC lysis buffer [10X], Biolegend, 420301) and dendritic cells were isolated using CD11c MicroBeads UltraPure Kit (Miltenyi Biotec, #130-125-835).

ELISA assays:

Total serum IgG (Mouse IgG ELISA kit, Cayman Chemicals, 501240), mouse IL2 (ELISA MAX Standard Set Mouse IL2, Biolegend, 431001) and OVA IgG1 (Anti-Ovalbumin IgG1(mouse) ELISA kit, Cayman, 500830) were measured using commercial kits.

To measure antibodies against corn, soy, wheat, or zein, high bind microwell 96 well plates (Corning, 9018) were coated with either the food of interest (corn/soy/wheat) smashed in PBS at 50 mg/ml in PBS or 5 mg/ml zein in 70% ethanol at 4°C overnight. The following morning, wells were washed 3X in PBS with 0.05% Tween-20 (Sigma) (PBST) and blocked for 2 hours at 37°C in 5% milk (Cell Signaling Technology) in PBS. Mouse serum samples were diluted 1:50 or 1:1000 (1:50 for tolerance contexts or zein, 1:1000 for allergy contexts). Serum samples were added to the wells and incubated for 1 hour at 37°C. Wells were washed 3x in PBST, then incubated in 1:2,500 anti-mouse (Promega, W4021) conjugated HRP antibody for 1 hour at 37°C. After 3 additional PBST washes, TMB reagent (Thermo Fisher, 50-493-593) was added and incubated until blue color developed, then stop solution (Biolegend, 423001) was added to quench the reaction. All ELISA data was collected on a Synergy HTX plate reader (Biotek) by measuring absorbance at 450 nm.

Co-culture experiment and ELISA assays:

50,000 DCs and 20,000 T cell hybridomas were co-cultured in 88% Dulbecco's Modified Eagle Media (DMEM, Fisher), 10% fetal bovine serum (Thermo Fisher), 1% penicillin-streptomycin (Thermo Fisher), 1% Glutamax (Gibco) for 24 hours with antigen formulation or peptides of interest. Supernatant was collected and used in an IL-2 ELISA assay (ELISA Max Standard Set Mouse IL-2, Biolegend, 431001) following manufacturer's instructions. Absorbance at 450 nm was measured using a Synergy HTX plate reader (Biotek). IL2 concentration was determined by generating a standard curve using log10 transformed concentration and absorbance values. In one experiment 20K hybridomas were plated in a 96-well U-bottom plate and co-cultured overnight with 50K FACS-purified CX3CR1+ macrophages (TCR β-/MHCIIint/F4/80+/ CX3CR1+/CD11c+) or dendritic cells (TCR β-/MHCIIhi/ F4/80-/CD11c+) from C57Bl/6 spleens in the presence of zein epitope. After overnight incubation, the supernatant was collected and assayed for IL-2 by ELISA (Biolegend) as described previously.

Bacterial screening of cDNA library:

An RNA sample was prepared from corn and soybean. The in-Fusion SMARTer Directional cDNA Library Construction Kit (Takara Bio USA) was used to generate a cDNA library with known DNA overhangs from 1 ug of input RNA. The resulting cDNA library was inserted into a plasmid for IPTG-inducible bacteria expression (pGEX-4T1, GE28-9545-49, Sigma-Aldrich) using Gibson Assembly (NEBuilder HiFi DNA Assembly Master Mix, NEB). Resulting plasmids were electroporated into ElectroMAX DH10B competent cells (18290015, Thermo Fisher). Bacteria were diluted and aliquoted into 96 well plates to yield approximately 30 colonies per well. The plates were recovered overnight. The following day, bacteria from each well was diluted into fresh LB media and the remainder was used to prepare glycerol stocks. The freshly diluted bacteria were incubated at 37°C with shaking until an OD of 0.5 was reached. IPTG (final concentration 0.5 mM Sigma-Aldrich) was added, and bacteria were incubated for an additional 16 hours at 18°C. Bacteria were then heat killed at 95 °C for 20 minutes and stored at -20 °C until use in co-culture experiments. Following the initial screen with this bacterial library, suspensions yielding signal on ELISA assay were recovered from glycerol stocks onto LB-agar plates with carbenicillin (Alfa Aesar). Individual colonies were selected and IPTG induced, as described for the library. Plasmids from bacteria that yielded ELISA signal were sequenced. This approach worked for corn and soybean but did not successfully identify an antigen for wheat.

Structure generation with Alpha-Fold and Pymol:

The structure of alpha-zein (uniprot ID: P02859), glycinin (uniprot ID: P04776), and gliadin (uniprot ID: A0A2R2Y406) were predicted using alpha-fold (*53*). The PDB file was downloaded and color coding for the image displayed was added in Chimera X (Version 1.7.1).

TCRdist:

TCR distance was calculated using the tcrdist3 python package (*54*). When a TRBV region returned two hits, only the first was included in the TCRdist calculation.

GLIPH searching:

The online graphical user interface version of GLIPH2 was used to find potential specificity groups.

*Hh* culture and oral infection

*Hh* was provided by J. Fox (MIT). *Hh* was cultured and passaged as previously described (*26*). Briefly, frozen aliquots of *Hh* were stored in Brucella broth with 20% glycerol and frozen at -80°C. *Hh* was cultured on blood agar plates (TSA with 5% sheep blood, Thermo Fisher). Inoculated plates were incubated in a hypoxia chamber at 37°C (Billups-Rothenberg). The hypoxia chamber was filled with an anaerobic gas mixture consisting of 80% nitrogen, 10% hydrogen, and 10% carbon dioxide (Airgas) to form a micro-aerobic atmosphere where oxygen concentration is approximately 3-5%. *Hh* cultured plates were incubated in hypoxia chambers for 4 days before animal inoculation. For oral infection, *Hh* was collected using a pre-moistened sterile cotton swab applicator tip to the colony surface and resuspended in Brucella broth. 0.3 mL of bacterial suspension administered to each mouse by oral gavage. Mice were inoculated with a second dose orally after 2 days. Confirmation of *Hh* colonization was performed using PCR analysis of fecal pellets.

Adoptive transfer of Hh7-2 TCR transgenic naïve CD4 T cells

Adoptive transfer of Hh7-2 tg naïve CD4 T cells were performed as previously described (*26*). Briefly, recipient mice were colonized with *Hh* via oral gavage seven days before adoptive transfer. Spleen and lymph nodes from donor Hh7-2 TCR transgenic mice were collected and mechanically dissociated through a 100 micron filter. Red blood cells were lysed using ACK lysis buffer (Lonza). Naïve Hh7-2 CD4 T cells were sorted as $CD4^+TCR\beta^+CD44^{lo}CD62L^{hi}CD25^-V\beta6^+$ on an Aria II (BD Biosciences). Cells were resuspended in PBS on ice and 50,000 cells were then transferred into congenic isotype-labelled recipient mice by retro-orbital injection. Cells from small intestine or colon lamina propria were analyzed 10-14 days after transfer.

RNA-sequencing:

Live/CD4+/CD25+/tetramer positive Tregs from chow fed mice, or Live/CD4+/CD25+/CD45.2+ cells from OVA fed mice were sorted on a BD Aria II. To achieve sufficient yields on the tetramer-specific and OVA-specific populations, samples from three mice were pooled together after staining and prior to flow sorting. Samples were sorted into Trizol reagent, frozen at -80, and sent to Genewiz/Azenta Life Sciences for Ultra-low input RNA-seq.

Total RNA was extracted from cells following the Trizol Reagent User Guide (Thermo Fisher Scientific). Extracted RNA samples were quantified using Qubit 2.0 Fluorometer (Life Technologies, Carlsbad, CA, USA) and RNA integrity was checked using Agilent TapeStation 4200 (Agilent Technologies, Palo Alto, CA, USA). Full-length cDNA synthesis and amplification was done using the SMARTSeq HT Ultra Low Input Kit (Clontech, Mountain View, CA), and the sequencing library was prepared with the Nextera XT DNA Library Preparation Kit (Illumina, San Diego, CA). The Agilent TapeStation was used for library validation, followed by quantification using the Qubit Fluorometer and quantitative PCR (KAPA Biosystems, Wilmington, MA). Sequencing was performed with the Illumina HiSeq instrument using a 2x150 Paired End configuration according to the manufacturer's instructions. The resulting sequence data (.bcl files) was converted into fastq files and de-multiplexed using bcl2fastq 2.17 Software (Illumina), allowing one mis-match for index sequence identification. Fastq files were quality checked using FastQC. Next, low-quality reads and adapter sequences were trimmed with Trimmomatic and FastQC was re-run to evaluate the quality. Reads were aligned to the mouse reference genome using STAR and resulting data were analyzed in DESeq2 (Version 1.42.0). For all heatmap visualizations and Chow vs AAD analysis (Differential gene table and volcano plot), the data were filtered with a 200 count minimum. For Zein vs Chow and Zein vs AAD analysis (Differential genes table and volcano plot) the minimum was increased to 4800 to narrow focus to differential genes expressed at a higher level. A heatmap to visualize the data was generated using PheatMap (Version 1.0.12). To make the heatmap, values of zero were replaced with 0.1 to allow for log scaling. Volcano plots were generated using the following packages: pandas (Version 1.5.3), matplotlib.pylab (Version 1.22.4), seaborn (Version 0.12.2) and numpy (Version 1.22.4). Violin plots were constructed using seaborn (Version 0.12.2).

Single cell transcriptome and TCR clonotype analysis:

Samples for single cell RNA-sequencing were categorized by bacterial colonization, as well as small versus large intestine. Cells from each animal were resuspended in 100 µl staining buffer (PBS with 4% FBS and 2mM EDTA) along with 0.2 micrograms of a unique hashing antibody to mark each category: 155861 TotalSeq C0301 anti-mouse Hashtag 1, 155863 TotalSeq C0302 anti-mouse Hashtag 1, 155865 TotalSeq™-C0303 anti-mouse Hashtag 3, 155867 TotalSeq™-C0304 anti-mouse Hashtag 4, 155869 TotalSeq™-C0305 anti-mouse Hashtag 5, 155871 TotalSeq™-C0306 anti-mouse Hashtag 6. Along with hashing, cells were stained with surface antibodies at 1:400 dilution, and appropriate tetramers at 1:100 dilution, followed by 3 washes with 200 µl staining buffer. Pooled cells were subsequently sorted on a FACSAriaII (BD Biosciences) using a 70-micron nozzle, operating FACSDiva software, with post-sort analysis performed on FlowJo 10.8.1 (Tree Star). Representative gating is demonstrated in Supplemental Figure 9. Approximately 66,000 total cells were collected into 5 ml complete media and maintained on ice. Viability and final cell counts were assessed with trypan blue and Countess II FL automated counter (ThermoFisher).

After passing through a 40-micron filter and resuspending in PBS, approximately 43,000 final cells were loaded onto three lanes of 10x Genomics Chip K, following manufacturer protocols. RNA, HTO, and TCR libraries were constructed strictly following the Chromium Next GEM Single Cell 5' dual index protocol (10X Genomics kit 1000265).
Single cell RNA-sequencing data were analyzed using Scanpy (version 1.9.4) (*55*). Data was filtered to only include: cells with a minimum of 500 genes, genes that were detected in a minimum of 10 cells, cells with fewer than 4500 genes, cells with fewer than 5% mitochondrial genes. Data were normalized: (1) for total count using the normalize_total function, (2) log transformed using the log1p function, (3) highly variable genes filtered with min_mean=0.0125, max_mean=10, min_disp=0.15, (4) regression on total counts and percent mitochondrial genes, (5) each gene scaled to unit variance with standard deviation clipped to 10. Next, data were reduced using principal component analysis and visualized as a umap projection using the leiden graph-clustering method. Tregs were identified as clusters of cells expressing Foxp3 and IL10. TCRs detected in the single cell sequencing were cloned onto hybridomas as described above. 10 TCRs were randomly selected from Tregs and 10 were selected from a cluster enriched in αZein$_{223-233}$ responsive Tregs. Gene expression scores were calculated using the Score_genes function in Scanpy.

<u>Treg activation methods:</u>
When indicated, single cell suspensions from lamina propria were incubated with 1X cell eBioscience™ Cell Stimulation Cocktail for 4 hours (ThermoFisher) or stimulated with 500,000 CD11c+ dendritic cells and 1 ug of peptide for 4 hours. To determine the potential impact of tetramer staining on phenotype, OT-II cells in one experiment were identified by both CD45.1/Cd45.2 staining and tetramer staining.

<u>Adoptive Treg transfer:</u>
Chow-fed mice were administered 10 ug αZein peptide or PBS emulsified in CFA. Ten days post-immunization, total CD4+CD25+ T cells from inguinal and axillary lymph nodes of were sorted, and ~$3\times10^5$ were transferred retro-orbitally into Zein-naïve CD45.1 mice born on AAD. The following day, recipient mice were then immunized with 10 αZein peptide emulsified in CFA, and inguinal and axillary lymph nodes were isolated and analyzed 10 days later.

<u>ELISpot Assay:</u>
Inflammatory response to *ex vivo* peptide restimulation was assayed by IFNg ELISPOT (Mouse IFN-γ Single-Color ELISPOT, CTL). $4\times10^5$ total lymphocytes were restimulated with 10ug/mL peptide in wells pre-coated with anti-IFNg capture antibody. After incubation at 37°C for 18 hr, plates were developed according to manufacturer instructions. Spots were quantified with an S6 Ultimate ELISPOT reader using ImmunoSpot ver7 software by CTL.

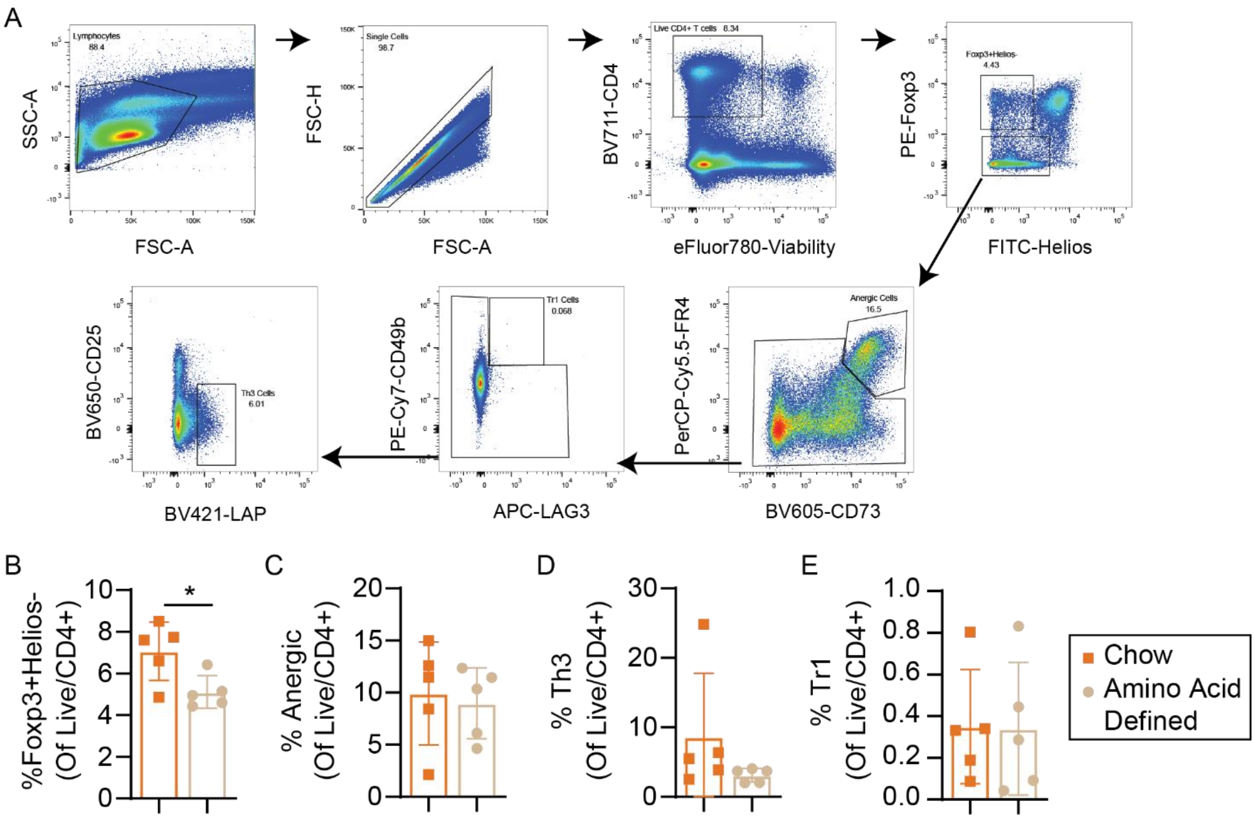


**Fig. S1. A protein containing diet induces Foxp3+/Helios- small intestine lamina propria Tregs.**

**(A)** Representative flow cytometry gating for identifying subsets of regulatory T cells. Percent **(B)** Foxp3+/Helios- peripheral Treg, **(C)** Anergic (folate receptor 4 [FR4]+/CD73+), **(D)** Th3 (CD25-/LAP+), and **(E)** Tr1 (CD49b+/LAG3+) cells in the small intestine lamina propria of mice fed amino acid defined or chow diets for 6 weeks starting just after weaning. N=5 mice/group. Data were generated in a single experiment. *P* values calculated using an unpaired t-test. Every dot represents an individual mouse. Error bars indicate mean ± SD. * denotes $p<0.05$, ** denotes $p<0.01$ and *** denotes $p<0.001$.

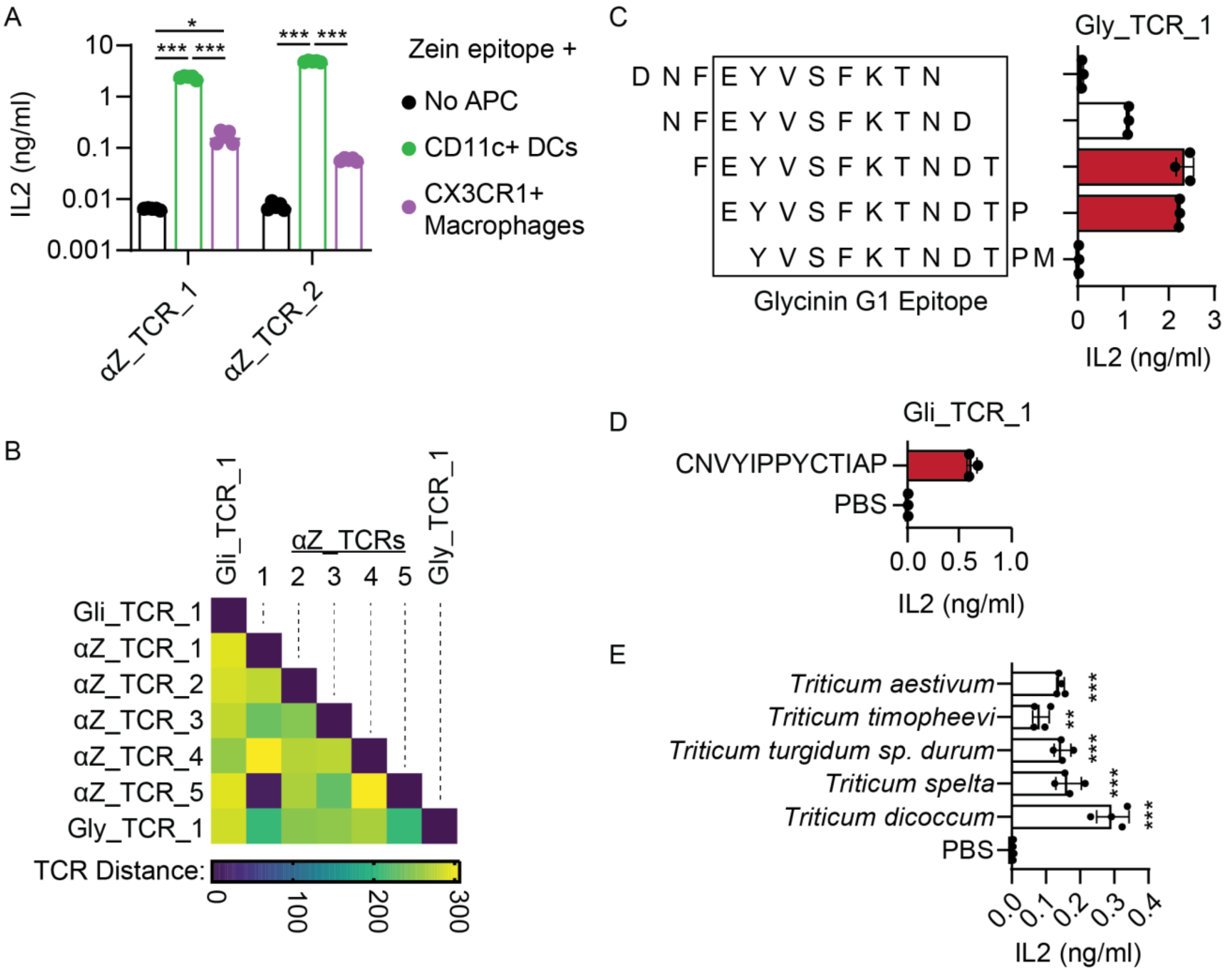


**Fig. S2. Receptor similarity and epitope mapping data for food-responsive TCRs.**

**(A)** TCR stimulation in hybridoma assay by CD11c+ dendritic cells compared to CX3CR1+ macrophages. All cultures were pulsed with zein epitope. **(B)** Similarity between TCRs was calculated using tcrdist3 (*1*). **(C)** Tiling scans with 10-11 amino acid peptides spanning glycinin G1 were used to identify minimum antigen epitope using hybridoma assays to measure TCR stimulation. **(D)** The gliadin-responsive TCR was screened with the candidate epitope CNVYIPPYCTIAP. Red bars do not differ significantly from each other ($p > 0.05$) and represent the maximum activation. **(E)** Gli_TCR_1 was stimulated with lysates from other wheat species reported to contain the same epitope (Table S4). N=4/group in panels A and E. N=3/group in panels C-D. Data were generated in a single experiment for Panels A, D, and E and are representative of two experiments in Panel D. *P* values were calculated using a two-factor ANOVA with Sidak's multiple comparisons test to compare effects of antigen presenting cell for each hybridoma (Panel A), a one-factor ANOVA with Dunnett's multiple comparisons test to compare each lysate against the PBS control (Panel C,E) or using an unpaired t-test (Panel D). Error bars indicate mean ± SD. Every dot represents a cell culture replicate. APC, antigen presenting cell. TCR, T cell receptor. * denotes $p<0.05$, ** denotes $p<0.01$ and *** denotes $p<0.001$.

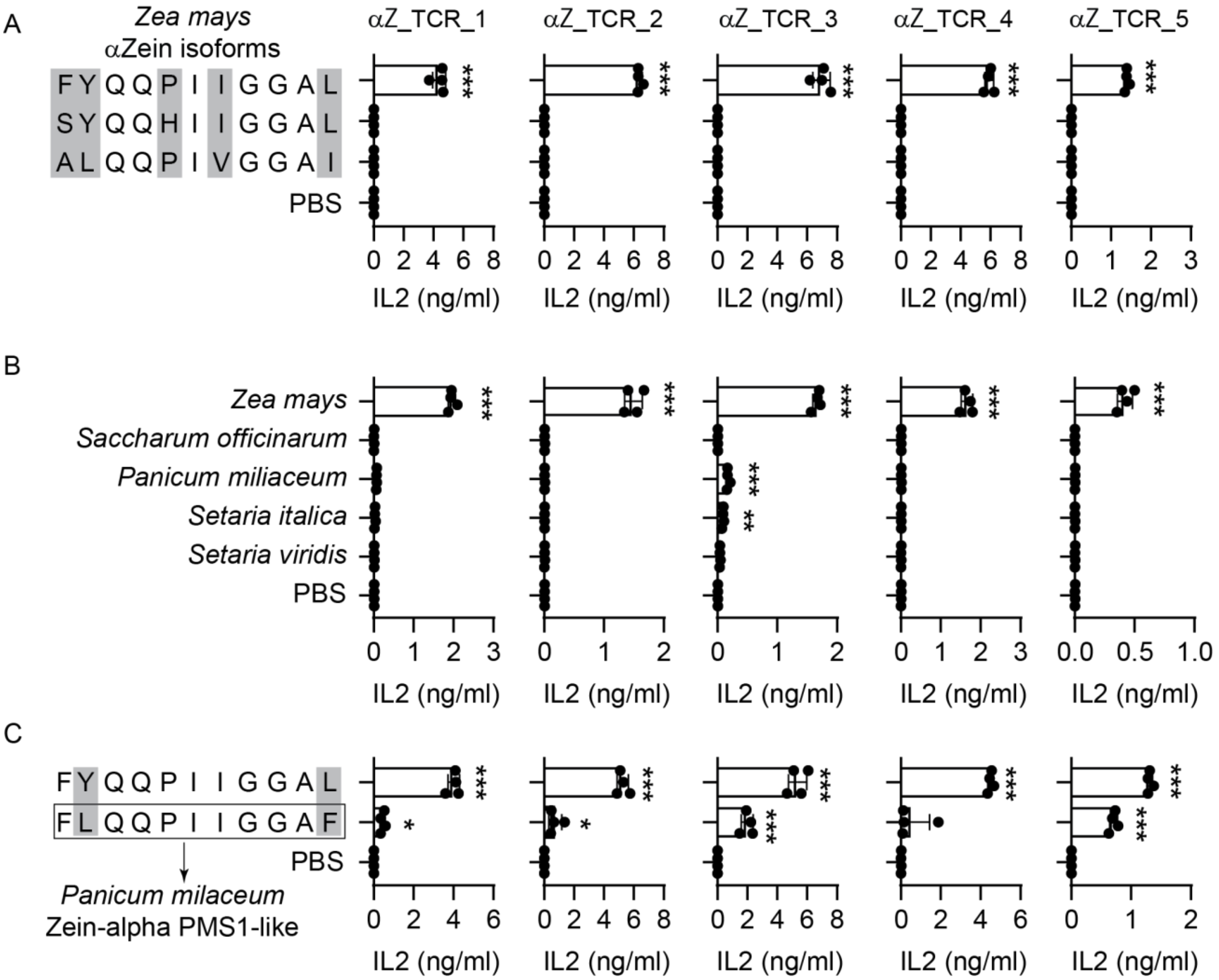


**Fig. S3. αZ_TCRs show minimal cross-reactivity toward other isoforms of zein, or potential homologs from other plants.**

Hybridomas expressing αZ_TCRs were stimulated using the mixed lymphocyte assay with **(A)** peptides representing three other 19 kDa alpha-zein sequence variants, **(B)** lysates from plants with a putative homologs, or **(C)** αZein$_{223\text{-}233}$ or a synthetic peptide from *Panicum milaecum.* N=4 replicates/condition (all panels). Data in Panels A and C were generated in a single experiment and data in Panel B are representative of 3 independent experiments. *P* values were calculated using 1-factor ANOVA with Sidak's multiple comparisons test to compare each lysate against the PBS control (in Panels A-C). Error bars indicate mean ± SD. Every dot represents a cell culture replicate. TCR, T cell receptor. * denotes $p<0.05$, ** denotes $p<0.01$ and *** denotes $p<0.001$ compared to PBS treatment.

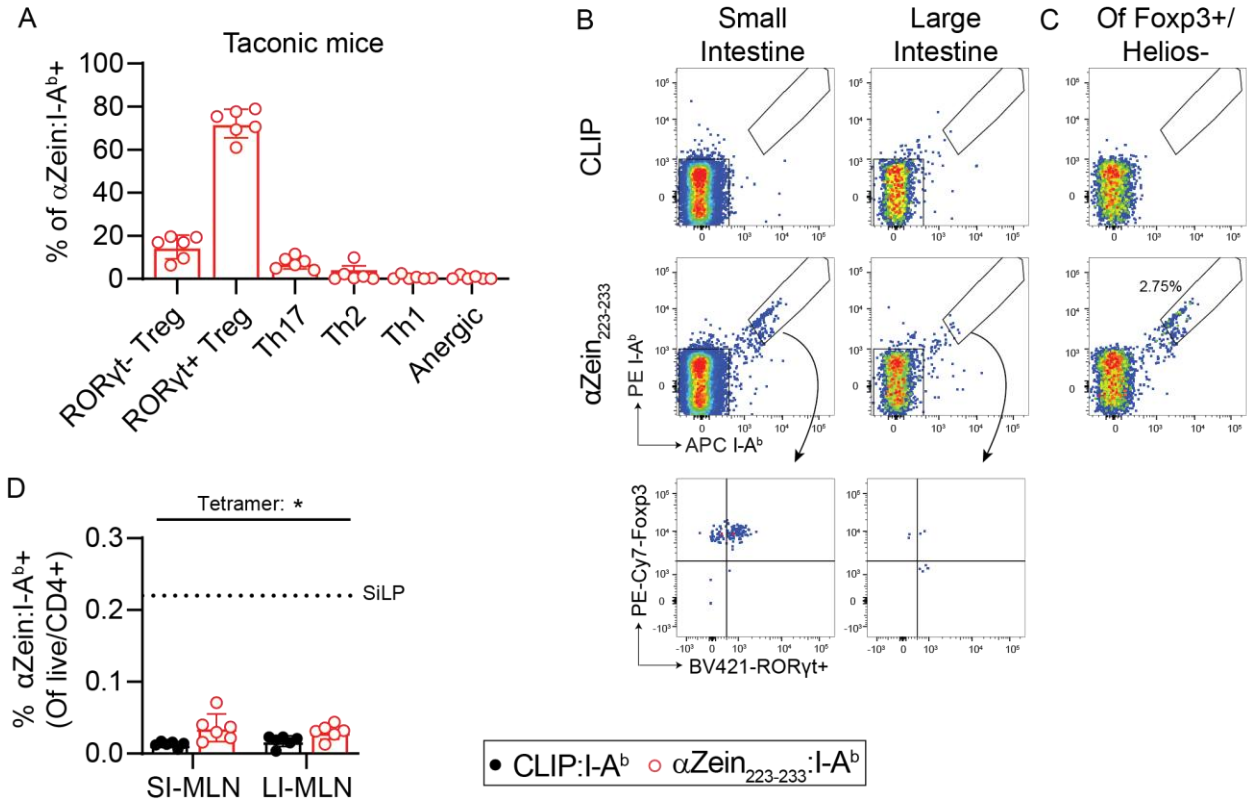


**Fig. S4. Characterization of αZein223-233 responsive T cells in small and large intestine**
(**A**) Distribution of αZein T cells in mice purchased from Taconic. **(B)** Abundance and phenotype distribution of αZein223-233 responsive T cells in small and large intestine in young adult mice (Pre-gated on Live/CD4+). Low numbers of Zein responsive cells in the large intestine preclude definitive subtype analysis. **(C)** Representative number of pTregs responsive to αZein223-233 (Pre-gated on Live/CD4+/Foxp3+/Helios-). **(D)** Tetramer positive cells in mesenteric lymph nodes draining the small and large intestine. The dotted line indicates frequency of tetramer positive cells observed in small intestine lamina propria (SiLP) sampled at the same time. N=6/group in Panels A and D. Data were generated in a single experiment. *P* values were calculated using a repeated-measured two-factor ANOVA (Panel D). Error bars indicate mean ± SD. Every dot represents an individual mouse. SI, small intestine. LI, large intestine. MLN, mesenteric lymph node. * denotes $p<0.05$, ** denotes $p<0.01$ and *** denotes $p<0.001$ compared to PBS treatment.

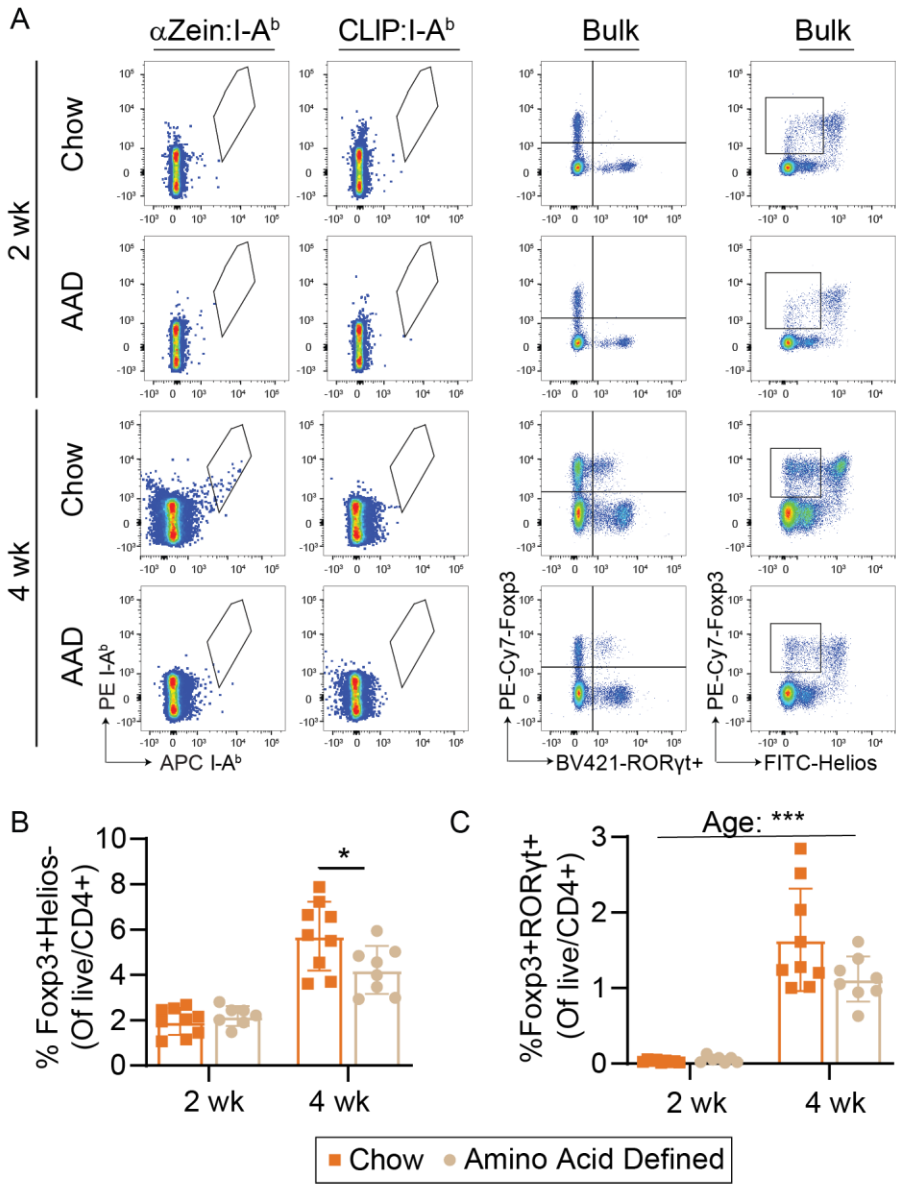


**Fig. S5. αZein$_{223-233}$ responsive T cells develop concurrent with weaning.**
Mice were born onto an amino acid defined (AAD) or chow diet and sacrificed at 2 or 4 weeks of age. **(A)** Flow cytometry gates related to figure 3D-. (Gated on live/CD4+) **(B)** Percent Foxp3+/Helios- or **(C)** Foxp3+/RORγt+ pTreg cells in the small intestine lamina propria (Gated on live/CD4+). N=7-9 mice/group/condition in panels B-C. In panel B data are representative of two experiments and the data in Panel C were generated in a single experiment. *P* values calculated by two-factor ANOVA with Sidak's multiple comparisons test when a significant interaction term was observed (Panel B-C). Error bars indicate mean ± SD. Every dot represents an individual mouse. * denotes p<0.05, ** denotes p<0.01 and *** denotes p<0.001 between indicated groups.

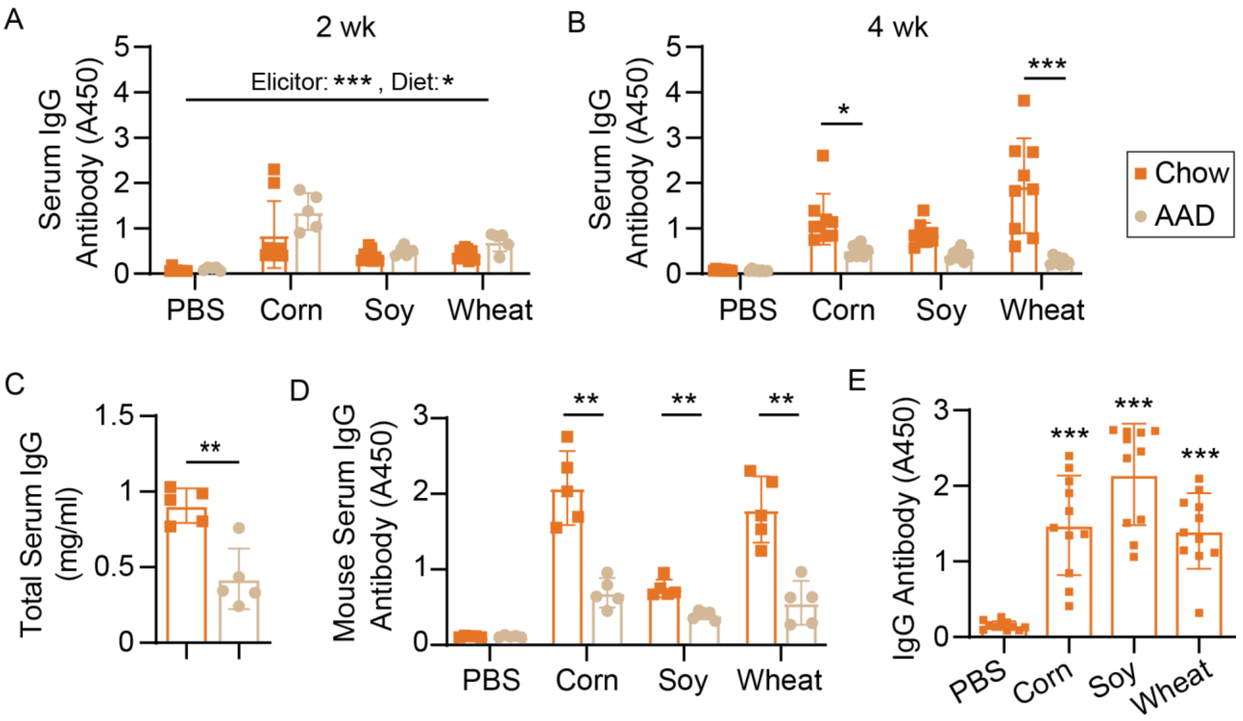


**Fig. S6. Development of food-responsive IgG antibodies during weaning.**
**(A-B)** Serum IgG antibodies targeting PBS (negative control), corn, soy, or wheat in mice born onto chow or AAD diets. In this experiment dams were randomized to diets at the end of pregnancy. **(C-D)** Total and food-specific IgG antibodies in mice consuming chow or AAD diets for 6 weeks starting at weaning. **(E)** Serum IgG antibodies targeting PBS, corn, soy, or wheat across a panel of human samples. N=7-9 mice/group/condition in panels A-B. N=5 mice/group in panels C-D. N=11 human donors in panel G. In Panels A-B, data are representative of two independent experiments, data in panel C-D were generated in a single experiment, and in Panel E the data are representative of the same assay performed twice on the serum samples. *P* values were calculated using a repeated measures two-factor ANOVA with Sidak's multiple comparisons test when a significant interaction term was observed (Panels A-B,D), unpaired t-test (Panel C), or repeated-measures one-factor ANOVA with a Dunnett's multiple comparisons test (Panel E). Error bars indicate mean ± SD. Every dot represents an individual mouse or human. AAD, amino acid defined. * denotes $p<0.05$, ** denotes $p<0.01$ and *** denotes $p<0.001$ between indicated groups, or for panel G, compared to the PBS control.

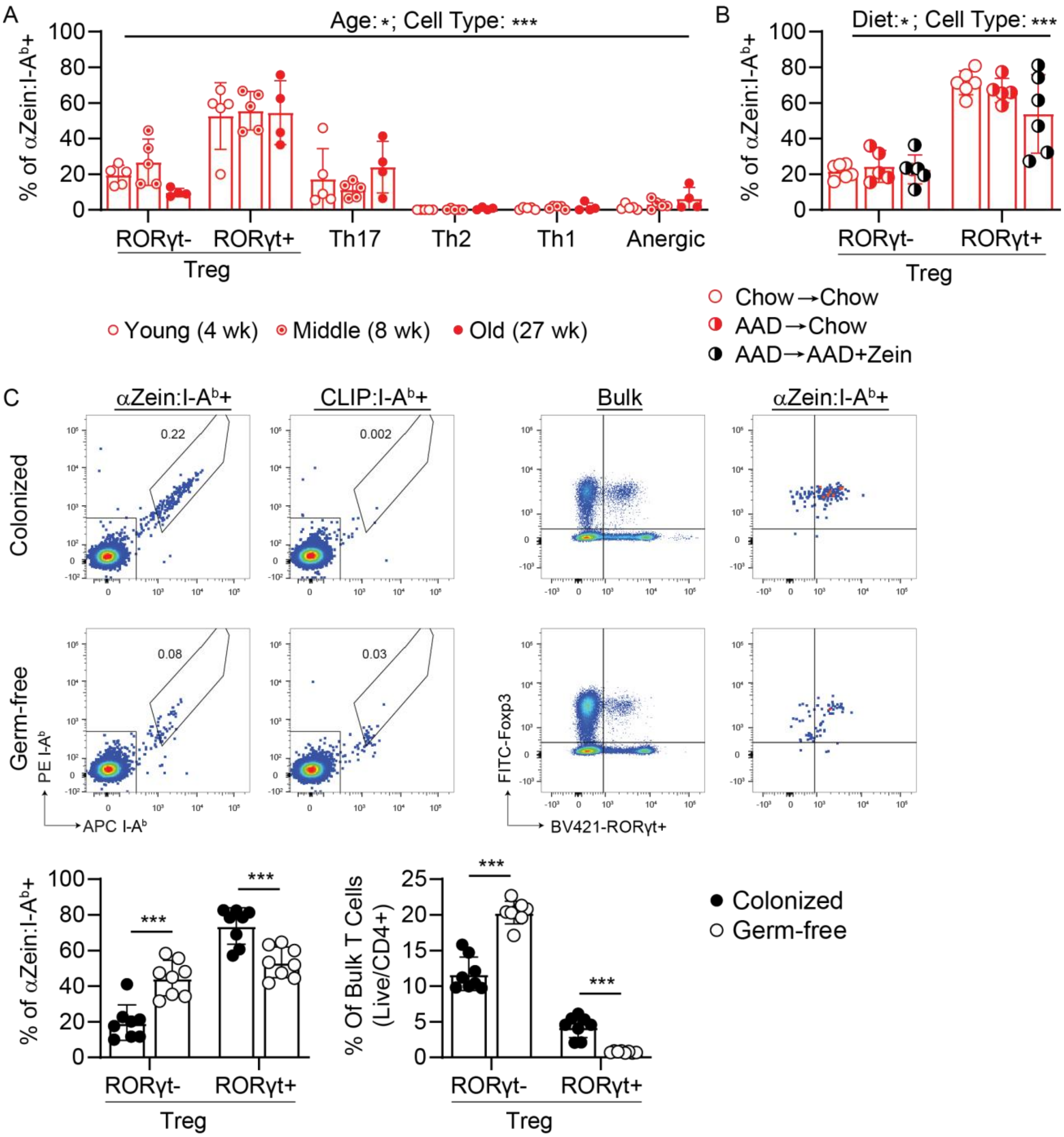


**Fig. S7. Zein T cell phenotype across age, diet, and microbiome status.**
**(A)** Phenotype distribution of αZein$_{223\text{-}233}$ responsive T cells across mice of different ages (Gated on Live/CD4+/ αZein:I-Ab-PE+/ αZein:I-Ab-APC+). **(B)** Phenotype distribution of αZein$_{223\text{-}233}$ responsive T cells across mice consuming chow, or who were swapped from AAD diet onto chow or AAD+10% zein from 6-8 weeks of age (Gated on Live/CD4+/ αZein:I-Ab-PE+/ αZein:I-Ab-APC+). **(C)** Phenotype distribution of αZein$_{223\text{-}233}$ responsive T cells and bulk T cells in colonized or germ free mice (Zein comparison gated on Live/CD4+/αZein:I-Ab-PE+/αZein:I-Ab-APC+ and bulk comparison gated on Live/CD4+). N=4-5/group in Panel A. N=5-6/group in Panel B. N=8/group in Panel C. Data were generated in a single experiment. *P* values were calculated using a two-factor ANOVA with Sidak's multiple comparisons test when

a significant interaction term was observed (Panels A-C). All experiments in this figure measure small intestine lamina propria samples. Error bars indicate mean ± SD. Every dot represents an individual mouse. AAD, amino acid defined. * denotes $p<0.05$, ** denotes $p<0.01$ and *** denotes $p<0.001$ between indicated groups.

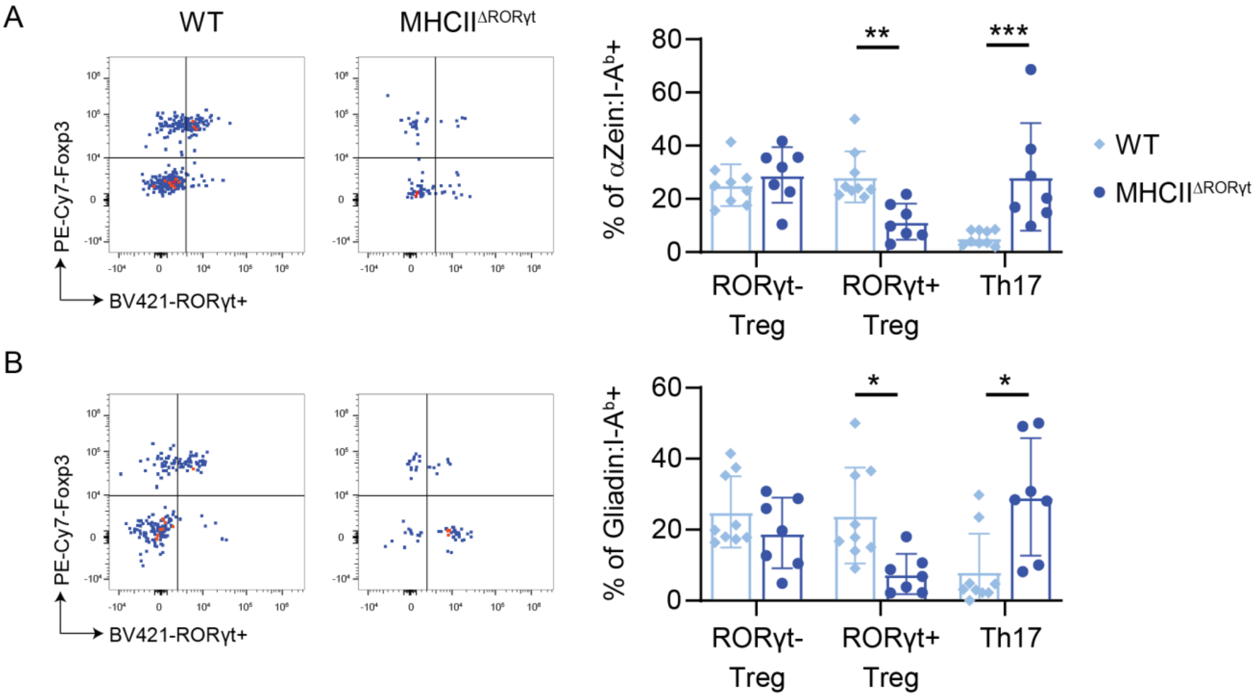


**Fig. S8. αZein223-233 and gliadin responsive T cells depend on RORγt+ antigen presenting cells for pTreg development.**

**(A)** αZein223-233 and **(B)** gliadin responsive cells were profiled for T cell subtypes in control mice (WT) or mice with an RORγt- driven MHCII deletion (MHCII$^{\Delta ROR\gamma t}$). Data were pre-gated on live/CD45+/TCRβ+/B200-/TCRγδ-/CD4+/αZein:I-Ab-PE+/αZein:I-Ab-APC+. N=7-9/group. Data are representative of two independent experiments. *P* values were calculated using a two-factor ANOVA. Error bars indicate mean ± SD. Every dot represents an individual mouse. * denotes $p<0.05$, ** denotes $p<0.01$ and *** denotes $p<0.001$.

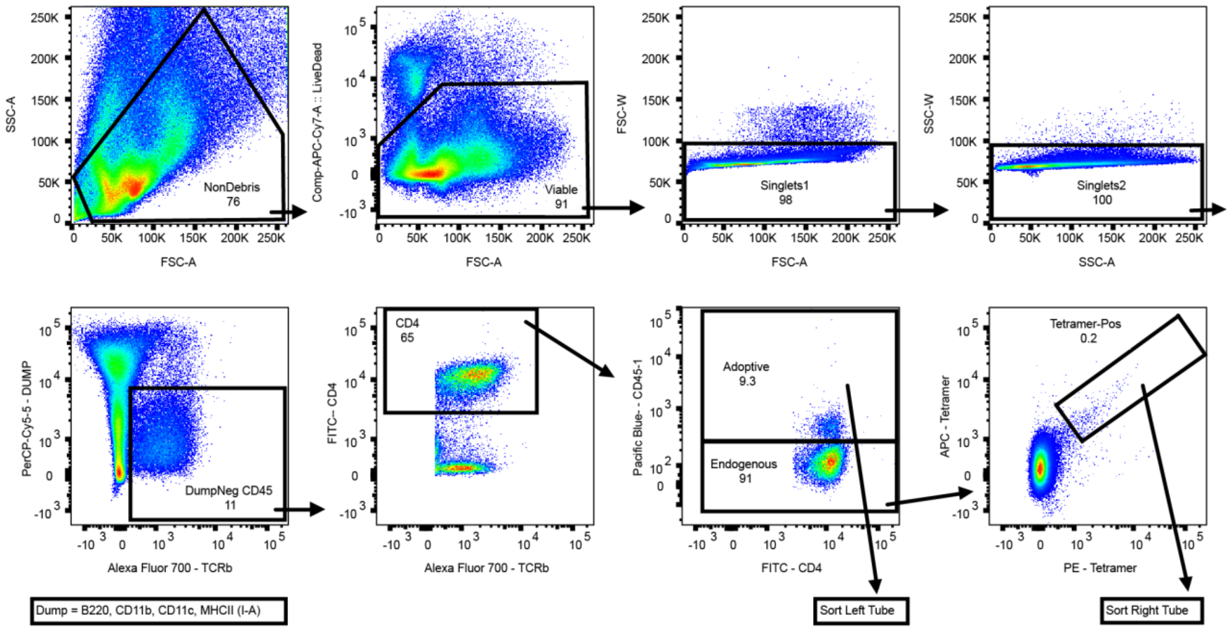


**Fig. S9. Gating strategy for single cell RNA-sequencing.**
Flow cytometry gates used for cell sorting for single cell RNA-sequencing.

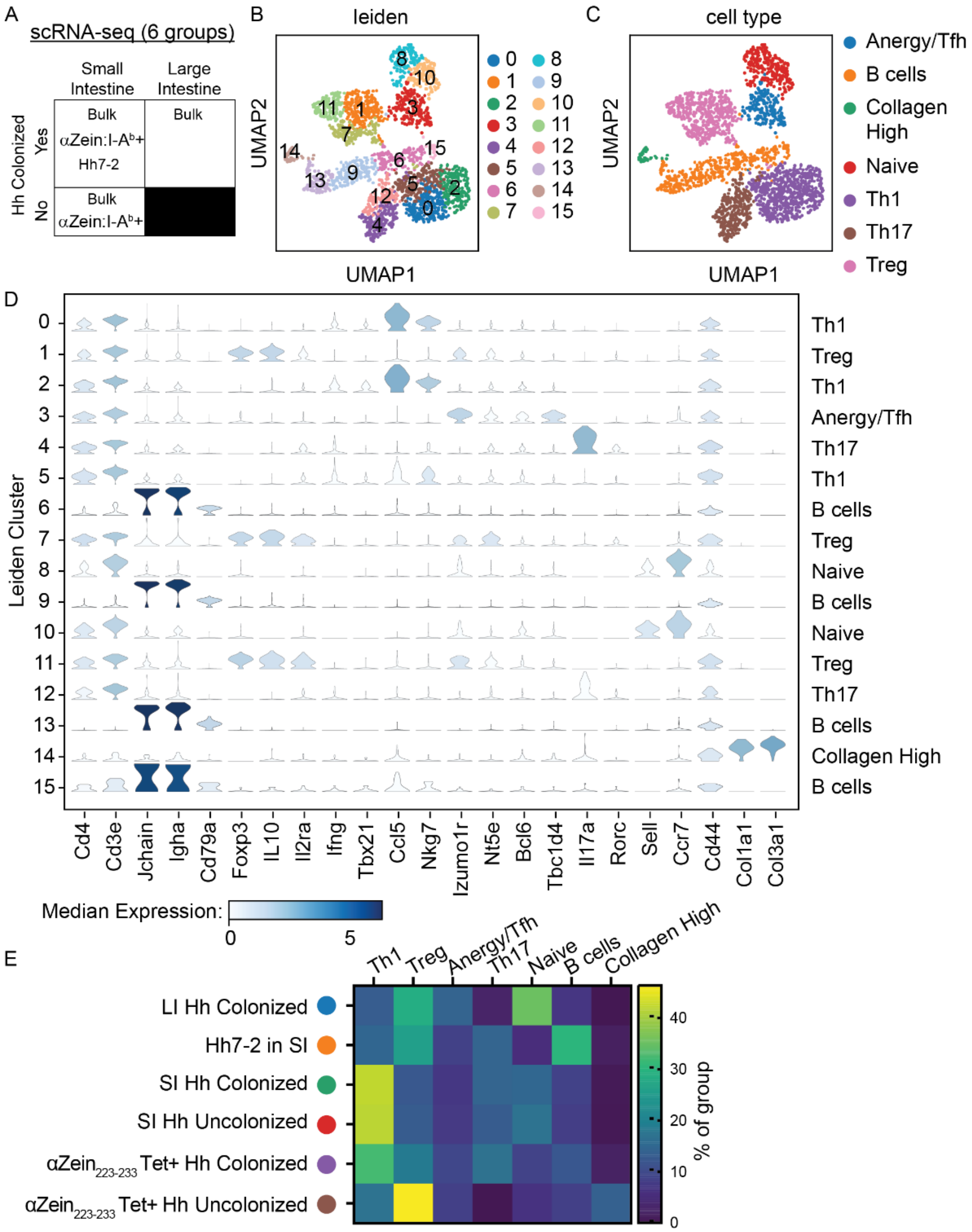


**Fig. S10. Clustering of single cell RNA-sequencing data into T cell types.**
**(A)** Table of sites selected for single cell RNA-sequencing of T cells: small intestine from *Hh* colonized or uncolonized mice, αZein$_{223\text{-}233}$ cells from Helicobacter hepaticus (*Hh)* colonized or uncolonized mice, adoptively transferred Hh7-2 cells, large intestine from *Hh* colonized mice. **(B-D)** The combined single cell transcriptomes grouped into 16 Leiden clusters that could be categorized into 7 major cell types **(E)** Distribution of cells with different (or unknown) antigen specificity across clusters. SI, small intestine.

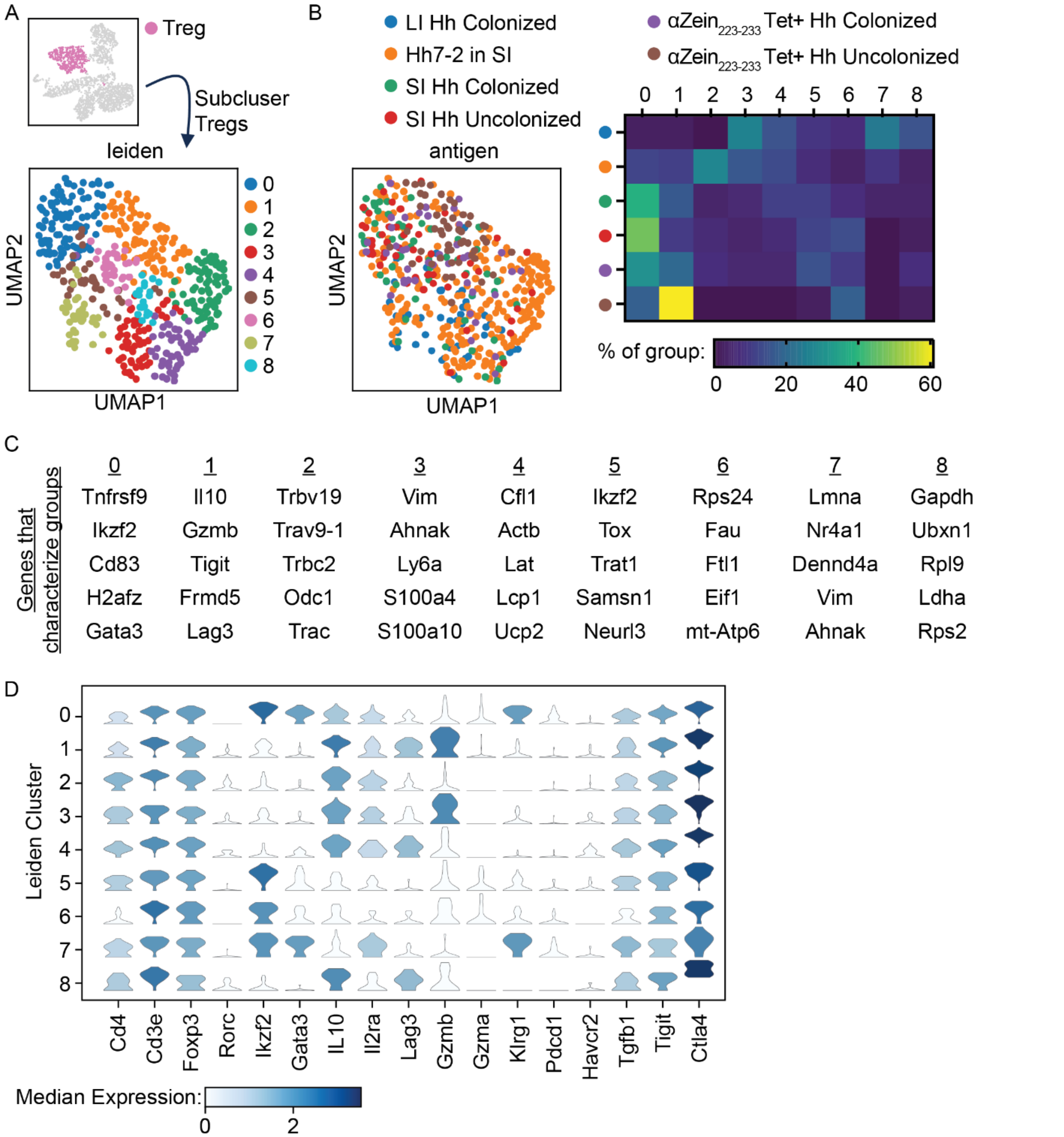


**Fig. S11. Immunosuppressive agents differentiate Tregs with different antigen specificities at the single cell level.**
**(A)** Leiden clustering of Tregs. **(B)** Distribution of cells across clusters. **(C)** Top 5 cluster defining transcripts identified by Scanpy. **(D)** Panel of Treg defining and immune suppressive markers across all Leiden clusters. Hh, Helicobacter hepaticus.

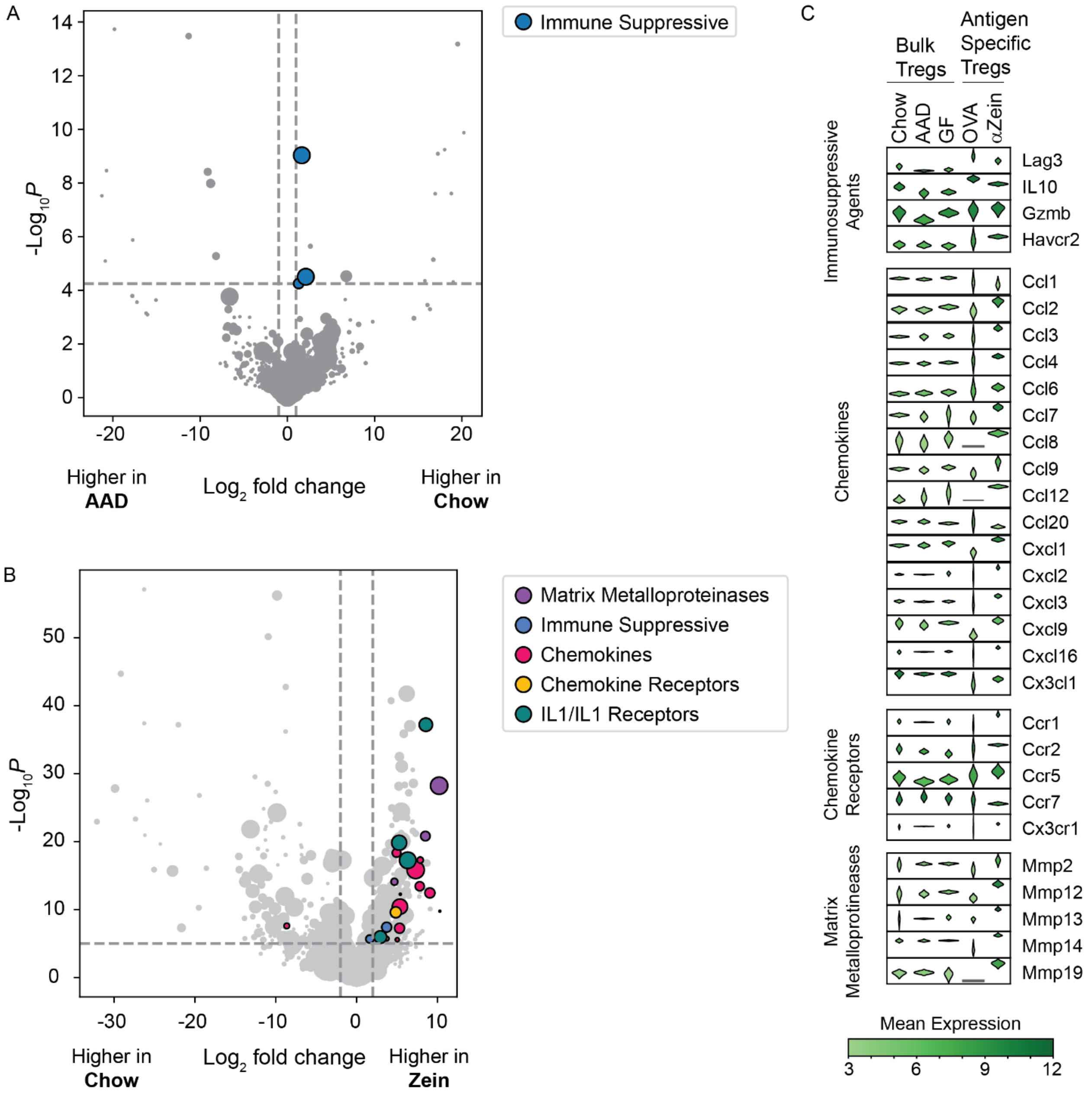


**Fig. S12. Identification of differential genes between Treg populations in bulk RNA-sequencing.**

Volcano plots showing differential genes between **(A)** Tregs from mice fed chow or amino acid defined diets or **(B)** αZein$_{223\text{-}233}$ Tregs compared to bulk Tregs from chow fed mice. Dot sizes are related to basemean of gene abundance with larger dots representing more abundant transcripts. **(C)** Violin plots of a subset of differential transcripts between Tregs from chow fed control mice, amino acid diet fed mice, germ-free mice, adoptively transferred OVA-specific Tregs or αZein$_{223\text{-}233}$ Tregs. GF, germ-free. AAD, amino acid defined.

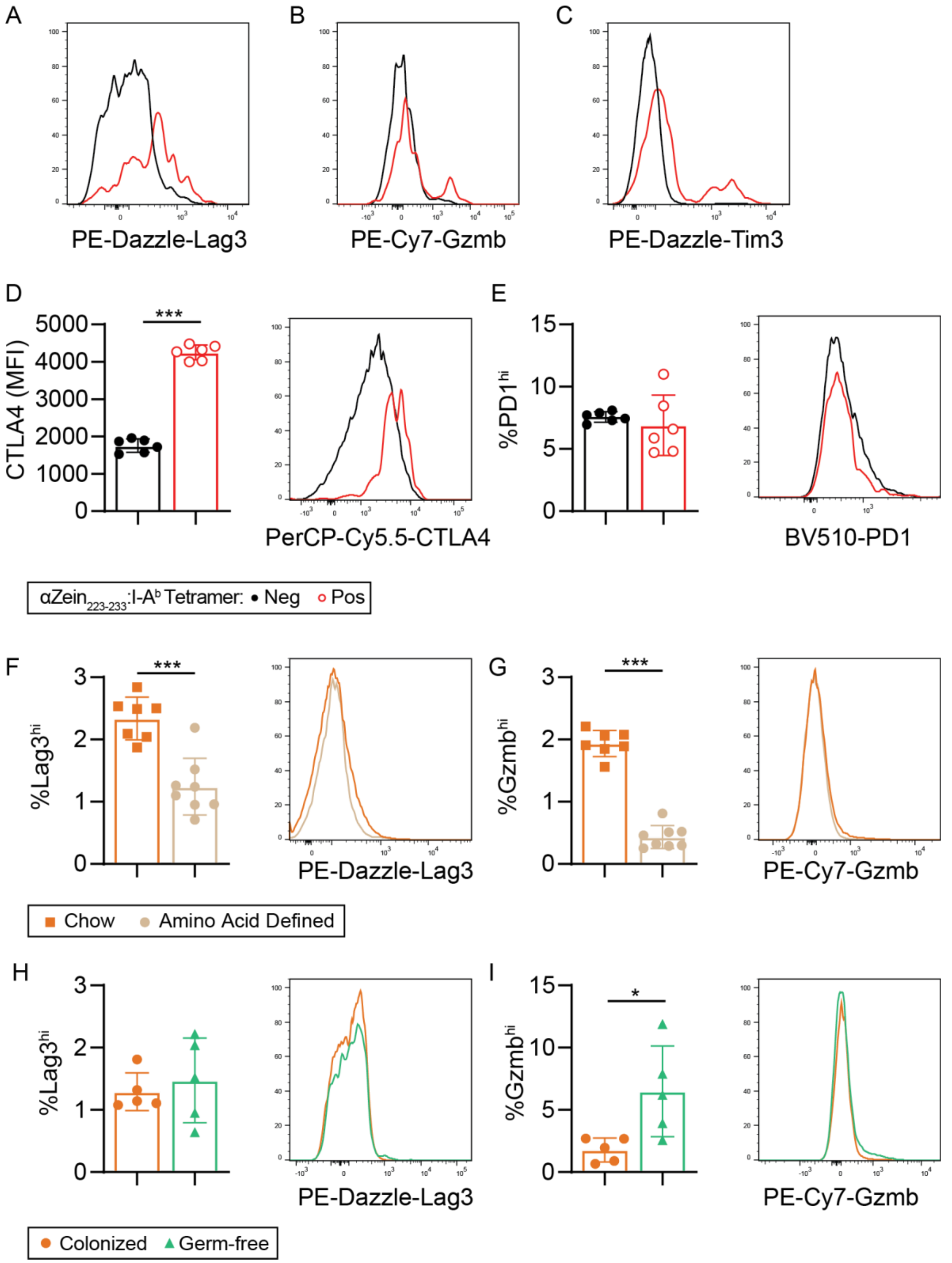


**Fig. S13. Immunosuppressive marker analysis and representative flow gating.**
**(A-C)** Flow cytometry gates related to Fig 4B-D. **(D-E)** CTLA4 MFI, and percent PD1$^{hi}$ cells between $\alpha$Zein$_{223\text{-}233}$ and tetramer negative Tregs. **(F-G)** Lag3 and Gzmb levels in Foxp3+ Tregs from chow or amino acid defined diet fed mice. **(H-I)** Lag3 and Gzmb levels between specific pathogen free and germ-free mice. All flow plots pre-gated on Live/CD4+/Foxp3+. N=6/group

in Panels D-E. N=7/group in Panels F-G. N=5/group in Panels H-I. *P* values calculated by paired (Panels D-E) or unpaired (Panel F-I) t-test. Data in panels D and G are representative of two independent experiments, data in Panel F is representative of 3 independent experiments, and data in panels E, H and I were generated in a single experiment. MFI, median fluorescence intensity. Every dot represents an individual mouse. * denotes p<0.05, ** denotes p<0.01 and *** denotes p<0.001.

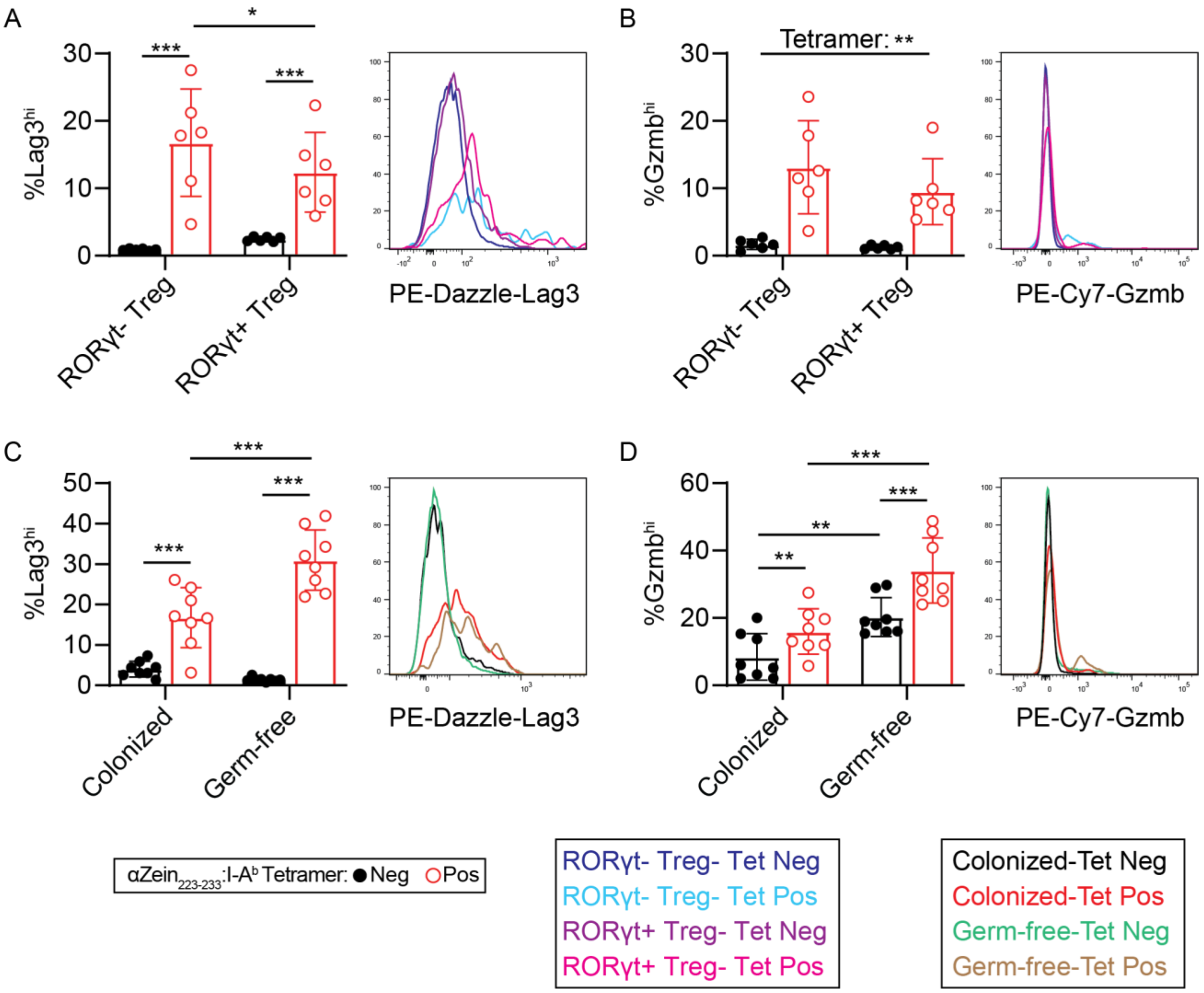


**Fig. S14. Lag3 and Gzmb levels in αZein Tregs based on RORγt expression and colonization status**

**(A-B)** Lag3 and Gzmb levels between $\alpha Zein_{223-233}$ binding and tetramer negative Tregs within categories of RORγt+ or RORγt- Tregs. **(C-D)** Lag3 and Gzmb levels between $\alpha Zein_{223-233}$ binding and tetramer negative Tregs between colonized and germ-free mice. N=6/group in Panels A-B and N=7/group in Panels C-D. All data were generated in a single experiment. *P* values calculated using a two-factor repeated measured ANOVA (Panels A-D) with an Uncorrected Fisher's LSD test (Panel A,C,D). Error bars indicate mean ± SD. Every dot represents an individual mouse. MFI, median fluorescence intensity. * denotes $p<0.05$, ** denotes $p<0.01$ and *** denotes $p<0.001$.

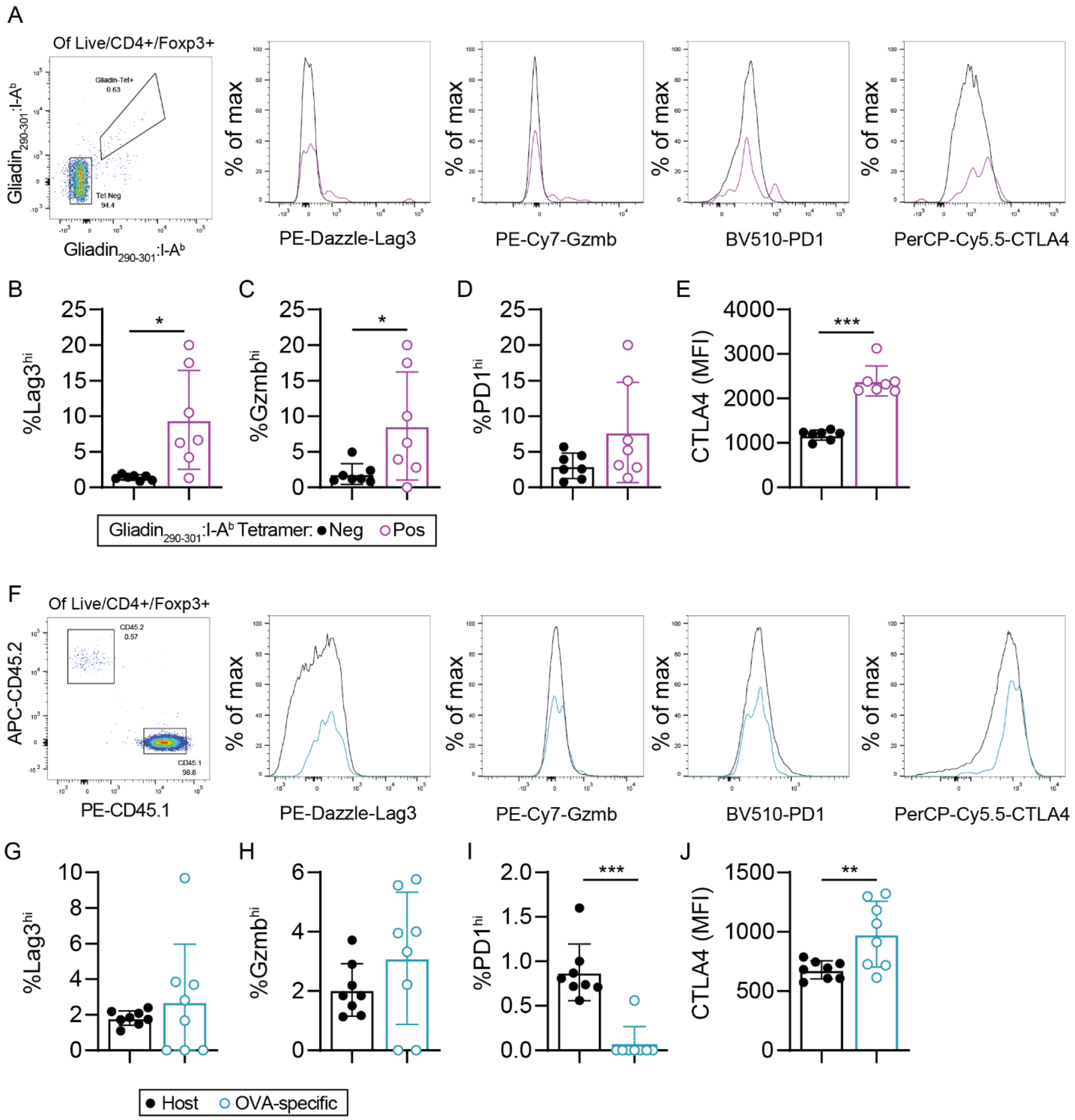


**Fig. S15. Select immunes suppressive markers in gliadin and OVA-specific Tregs.**
**(A-E)** Lag3, Gzmb, PD1, and CTLA4 levels between gliadin$_{273-285}$ and tetramer negative Tregs. **(F-J)** Lag3, Gzmb, PD1, and CTLA4 levels between adoptively transferred OT-II Tregs and host Tregs. Pre-gated on Live/CD4+/Foxp3. N=7-8/group. All data are representative of two independent experiments. *P* values calculated by paired t-test. Error bars indicate mean ± SD. Every dot represents an individual mouse. MFI, median fluorescence intensity. * denotes p<0.05, ** denotes p<0.01 and *** denotes p<0.001.

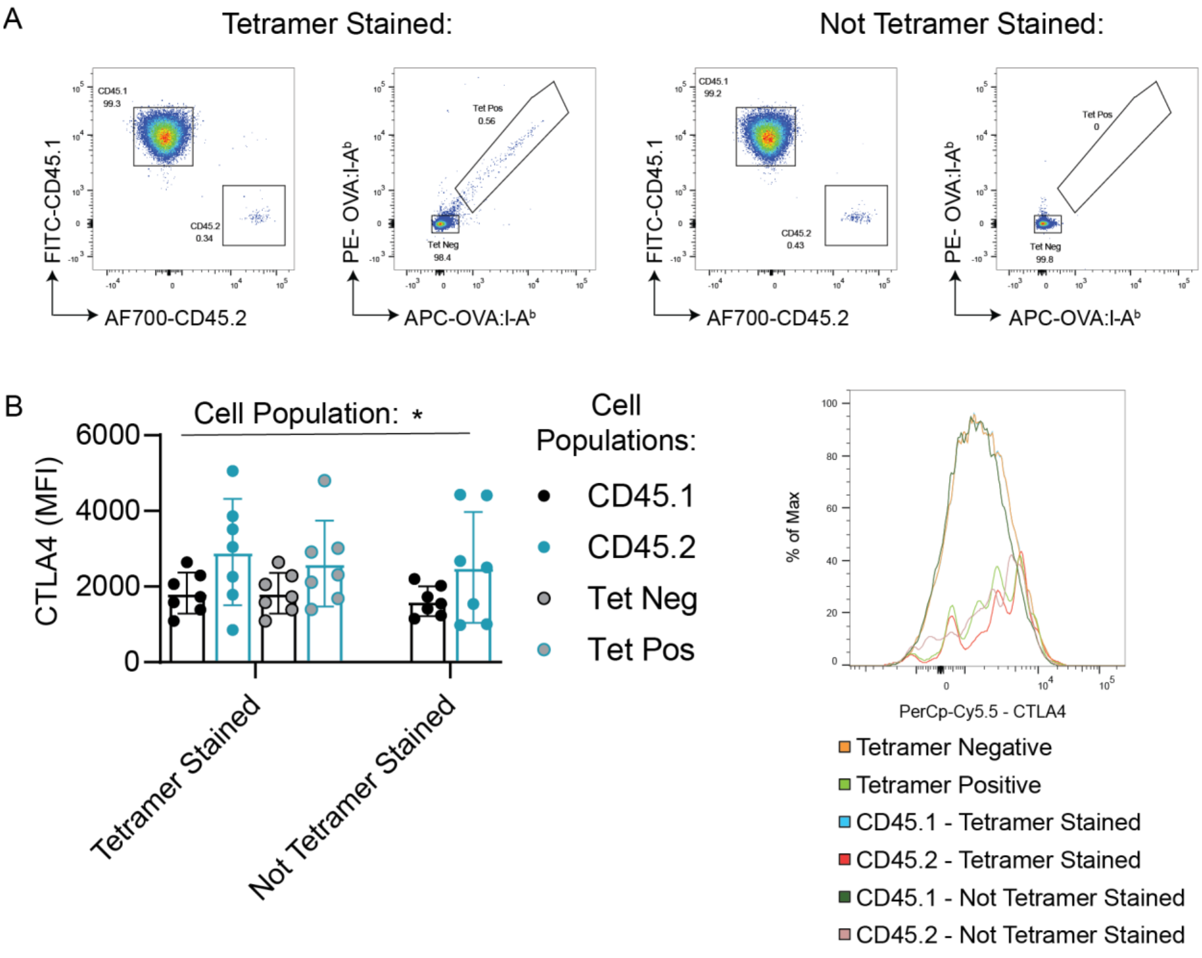


**Fig. S16. Tetramer staining does not alter cell profile of OVA-responsive Tregs.**
**(A)** CD45.2+ OTII cells were adoptively transferred into CD45.1 hosts. After 1 week of oral OVA exposure, lamina propria T cells were harvested. Samples were divided into tetramer stained and unstained conditions and additionally, OVA-responsive cells were identifiable by the CD45.1/CD45.2 markers. CTLA4 was used as the readout to check for potential effect of tetramer staining on cell phenotype. **(B)** CTLA4 in OVA-specific or host Tregs identified by CD45.1/CD45.2 staining or identified by tetramer staining. Data pre-gated on live/CD4+/Foxp3+. N=7/group. Data were generated in a single experiment. *P* values calculated by two-factor ANOVA. Error bars indicate mean ± SD. Every dot represents an individual mouse. * denotes $p<0.05$, ** denotes $p<0.01$ and *** denotes $p<0.001$.

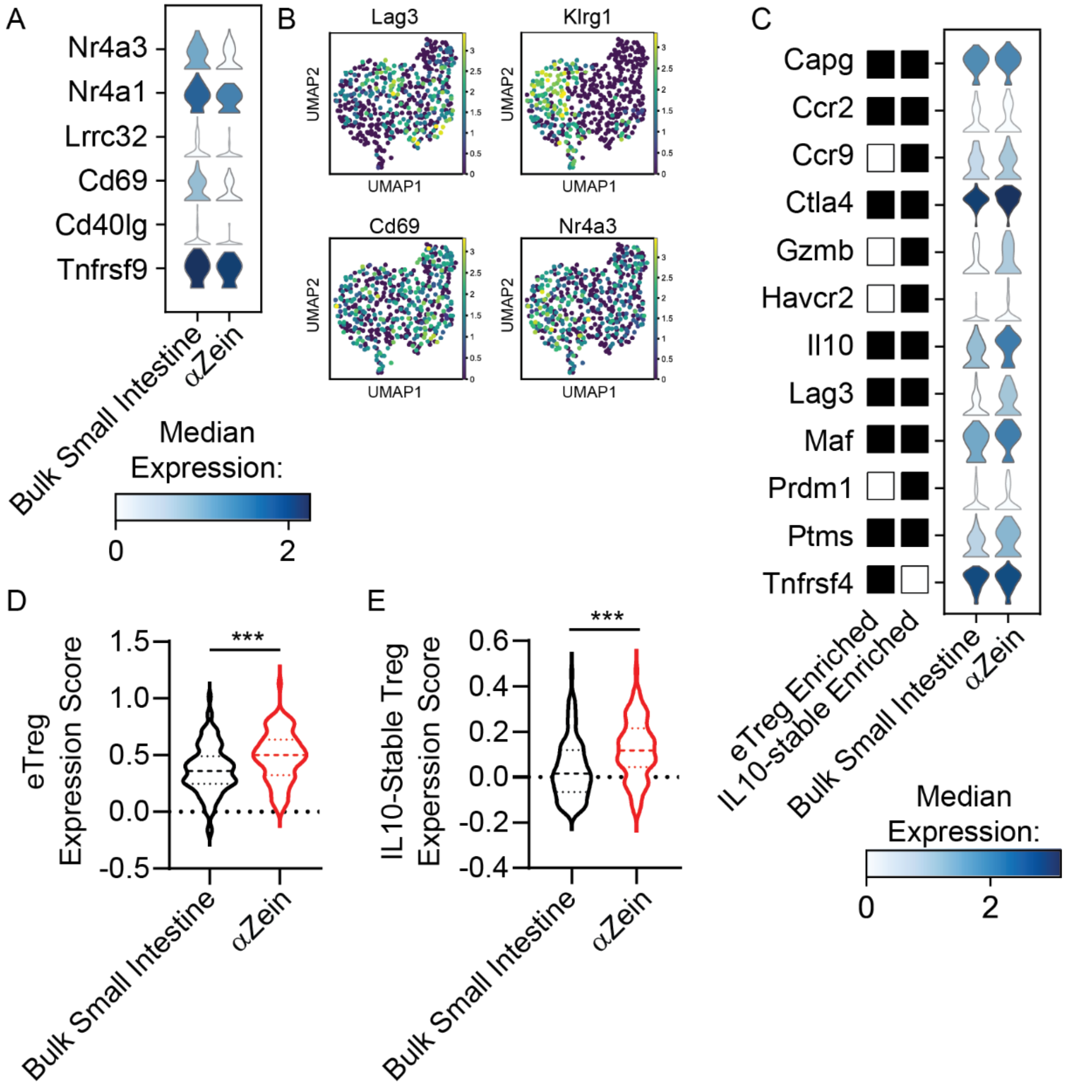


**Fig. S17. αZein Tregs show a gene expression profile similar to effector Tregs.**
**(A)** Single cell RNA-seq data showing canonical activation markers in bulk small intestine and αZein responsive Tregs. **(B)** UMAP plots from the single cell RNA-seq data showing Lag3, Klrg1, Cd69, and Nr4a3 across all profiled Tregs. **(C)** Levels of select differential genes that define eTreg or IL10-stable Treg populations measured in bulk small intestine Tregs or αZein-specific Tregs. **(D-E)** Single cell RNA-seq data of bulk small intestine Tregs or αZein Tregs were scored for similarity to published gene expression profiles representative of eTregs or IL10-stable Tregs. *P* values were calculated by unpaired t-test. *** denotes $p<0.001$.

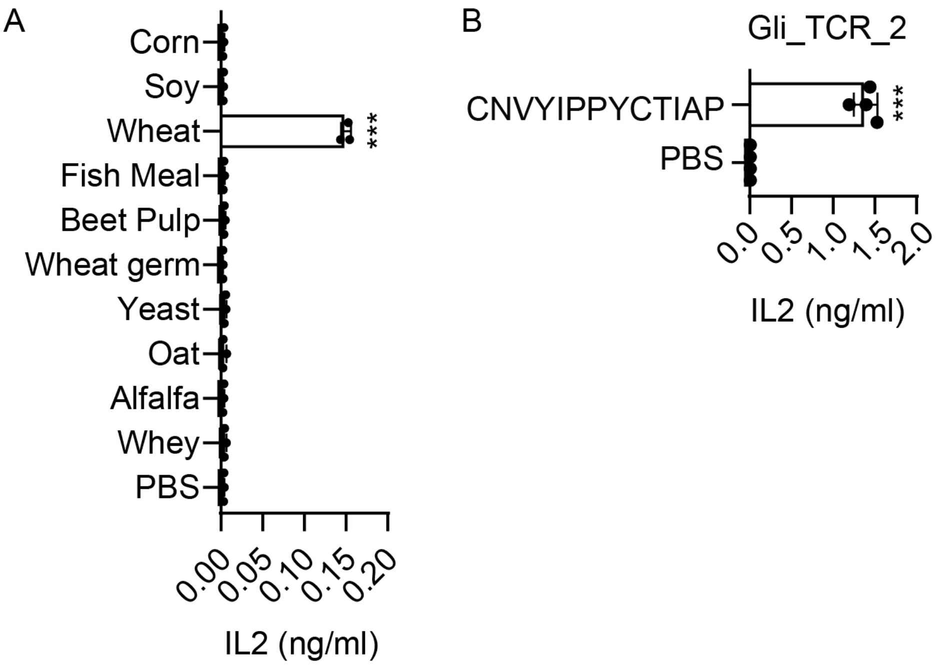


**Fig. S18. Identification of an additional gliadin-responsive TCR.**
**(A)** Identification of a wheat-responsive TCR (Gli_TCR_2). **(B)** Mapping of Gli_TCR_2 to an epitope. N=3-4/group. Error bars indicate mean ± SD. Every dot represents a cell culture replicate. TCR, T cell receptor. Data in panel A are representative of two independent experiments and in panel B were generated in a single experiment. *P* values were calculated by one factor ANOVA (Panel A) or unpaired t-test (Panel B). * denotes p<0.05, ** denotes p<0.01 and *** denotes p<0.001 compared to the PBS control.

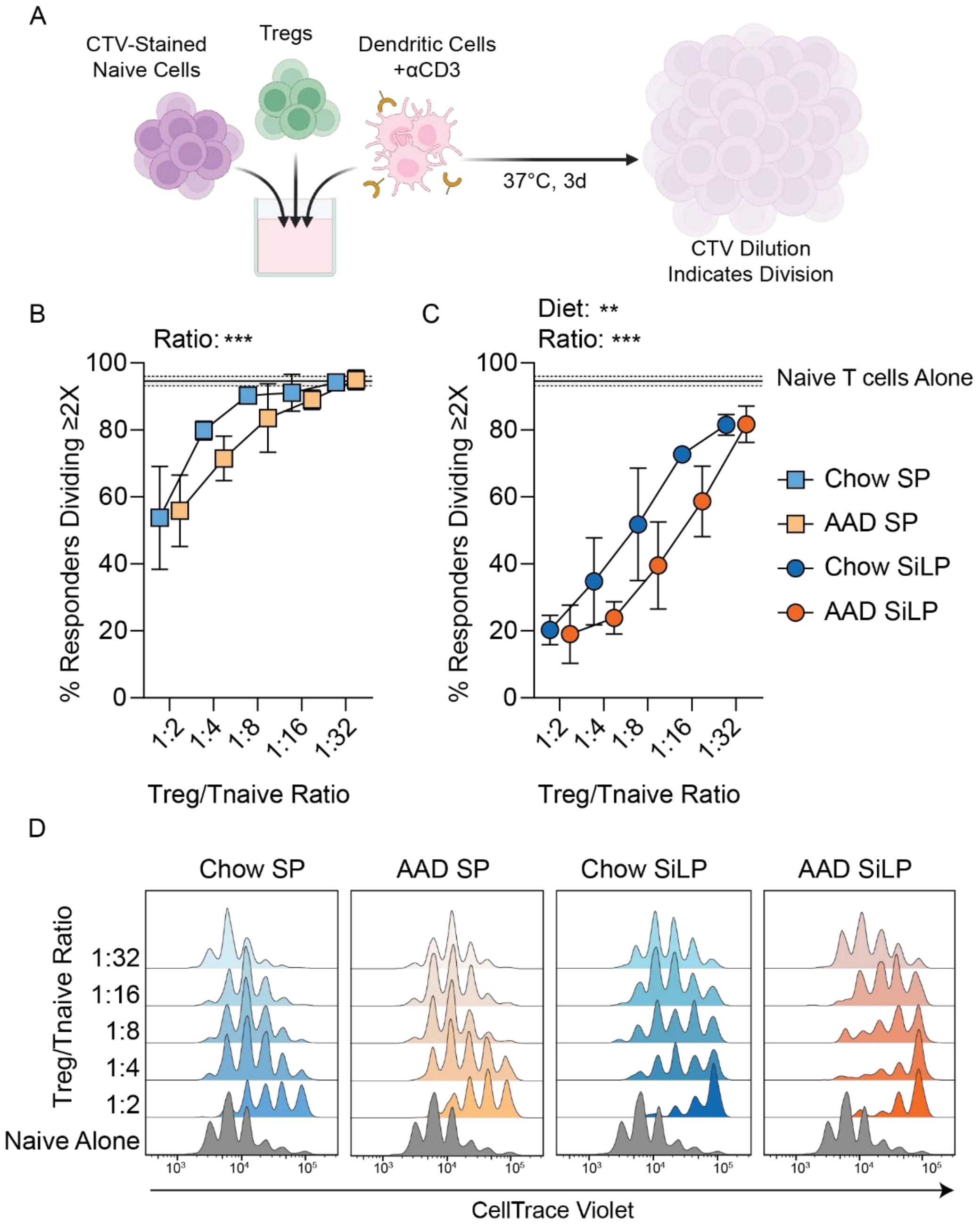


**Fig. S19. Suppressive capacity of Tregs from different anatomical sites in mice consuming Chow or AAD diets toward naïve T cells in an ex vivo assay.**
(**A**) Suppressive capacity was measured by monitoring CellTrace Violet dye dilution in naïve T cells indicating cell division after a 3 day co-culture with variable numbers of Tregs and dendritic cells and αCD3 antibodies to facilitate activation. **(B-D)** Naïve T cell division in bulk Tregs from small intestine lamina propria (SiLP) or spleen (SP) from mice born on chow or amino acid defined (AAD) diets. N=4-5/group. Data were generated in a single experiment. *P* values were calculated using a two-factor repeated measures ANOVA (Panel B-C). Dots represent the average of data from Tregs from 4-5 mice (Tregs from each animal cultured

independently). Error bars indicate mean ± SD. CTV, CellTrace Violet. * denotes $p<0.05$, ** denotes $p<0.01$ and *** denotes $p<0.001$ for the indicated factor.

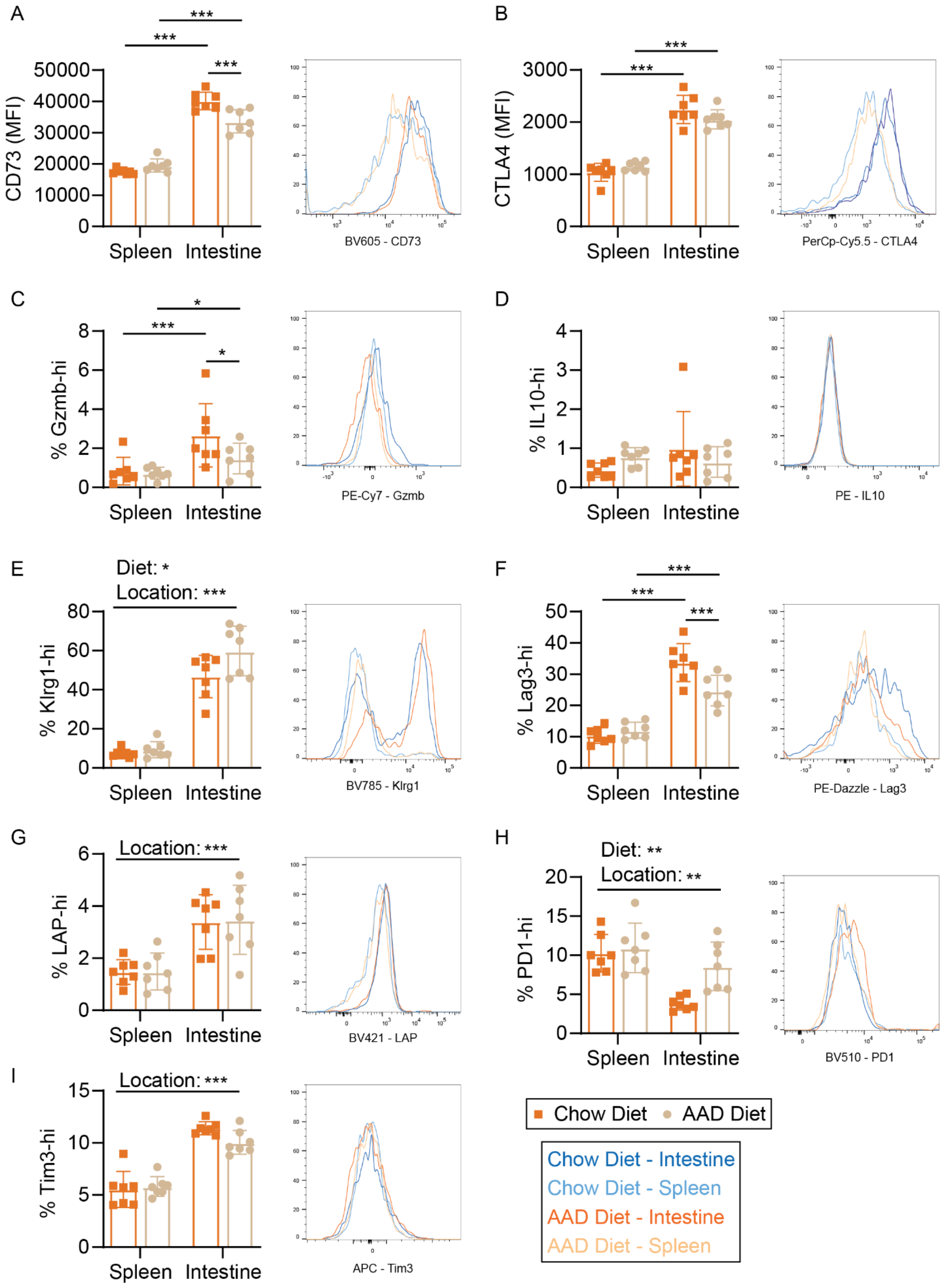
A
CD73 (MFI)
Spleen Intestine
BV605 - CD73
B
CTLA4 (MFI)
PerCp-Cy5.5 - CTLA4
C
% Gzmb-hi
PE-Cy7 - Gzmb
D
% IL10-hi
PE - IL10
E
Diet: *
Location: ***
% Klrg1-hi
BV785 - Klrg1
F
% Lag3-hi
PE-Dazzle - Lag3
G
Location: ***
% LAP-hi
BV421 - LAP
H
Diet: **
Location: **
% PD1-hi
BV510 - PD1
I
Location: ***
% Tim3-hi
APC - Tim3
Chow Diet AAD Diet
Chow Diet - Intestine
Chow Diet - Spleen
AAD Diet - Intestine
AAD Diet - Spleen

**Fig. S20. Immune suppression profile of Tregs from different anatomical sites in mice consuming Chow or AAD diets following ex vivo anti-CD3 activation.**
**(A-I)** CD73, CTLA4, Gzmb, Il10, Klrg1, Lag3, LAP, PD1, and Tim3 levels in Tregs from small intestine lamina propria or spleen samples from mice born onto chow or born onto amino acid defined (AAD) diets. N=7/group. Data were generated in a single experiment. *P* values calculated by two-factor ANOVA with an Uncorrected Fisher's LSD multiple comparisons test when a significant interaction term was observed. Error bars indicate mean ± SD. Every dot represents an individual mouse. MFI, median fluorescence intensity. * denotes $p<0.05$, ** denotes $p<0.01$ and *** denotes $p<0.001$.

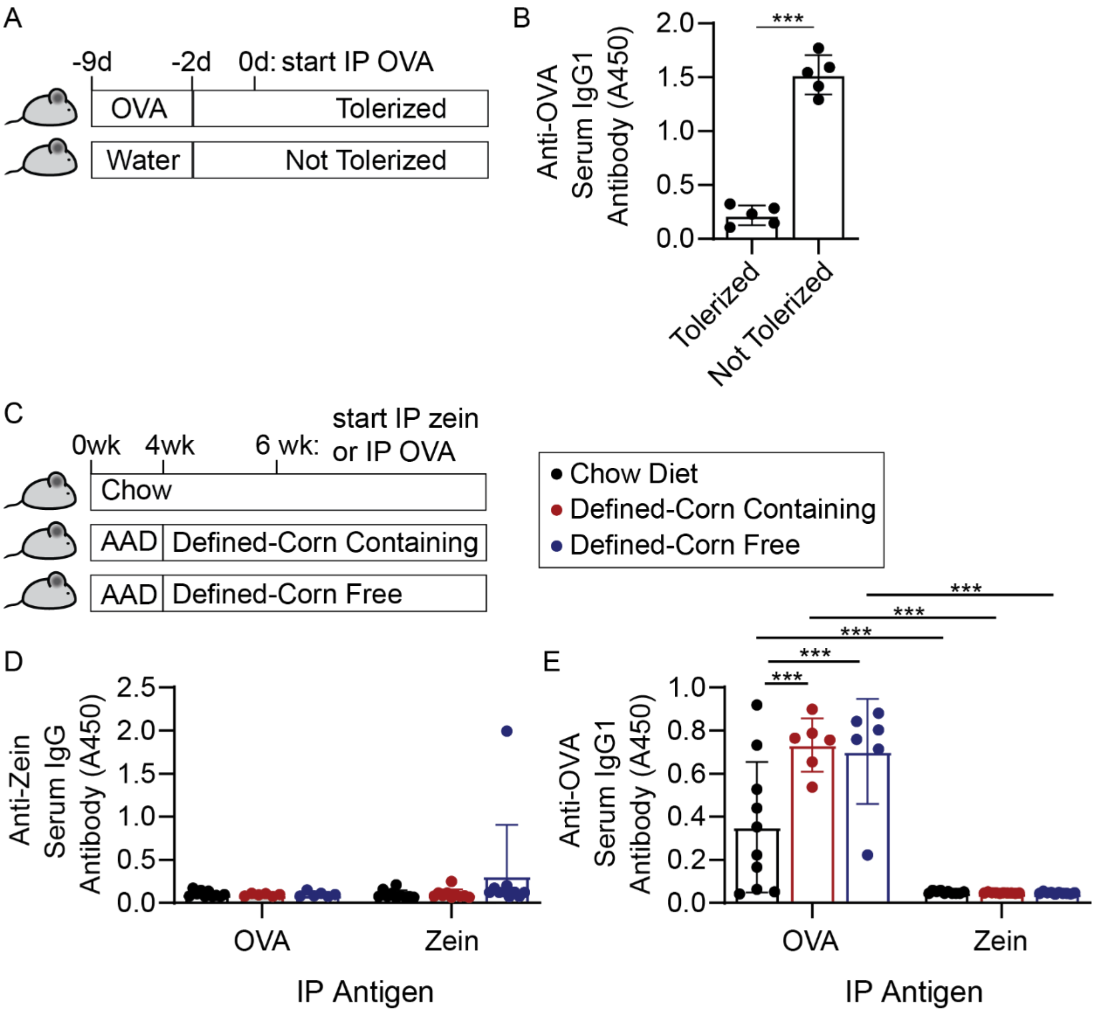


**Fig. S21. Tolerance to dietary proteins depends on diet**
**(A)** To induce OVA tolerance mice were given 10 mg/ml OVA in drinking water for 1 week or water alone as a control. On day 0 mice received an IP injection of OVA in alum and 14 days later mice received a booster injection with OVA alone. On day 21, mice were sacrificed and serum was collected. **(B)** Consuming OVA in drinking water induces oral tolerance reflected as lower anti-OVA IgG1. **(C)** Control mice always consumed chow. Experimental mice were born onto AAD diet then randomized at 4 weeks to consume a defined corn-containing diet (AAD + 10% corn + 10% wheat + 10% oat + 10% soy) or defined corn-free diet (AAD + 10% wheat + 10% oat + 10% soy). At 6 weeks of age, all mice received an IP injection of either zein or ova in alum and 14 days later received a booster injection of antigen alone. At 21 days after the first IP injection mice were sacrificed and serum was collected. **(D)** Anti-zein IgG antibodies in serum of mice described in panel D. **(E)** Anti-OVA IgG antibodies in serum of mice described in panel D. N=5-10 mice/group. Data in panel B is representative of two independent experiments and data in panels D-E were generated in a single experiment. *P* values calculated by unpaired t-test (Panel B), two-factor ANOVA (Panel D), or a two-factor ANOVA with Tukey's multiple comparisons test (Panel E). Error bars indicate mean ± SD. Every dot represents an individual mouse. AAD, amino acid defined. IP, intraperitoneal. * denotes p<0.05, ** denotes p<0.01 and *** denotes p<0.001.

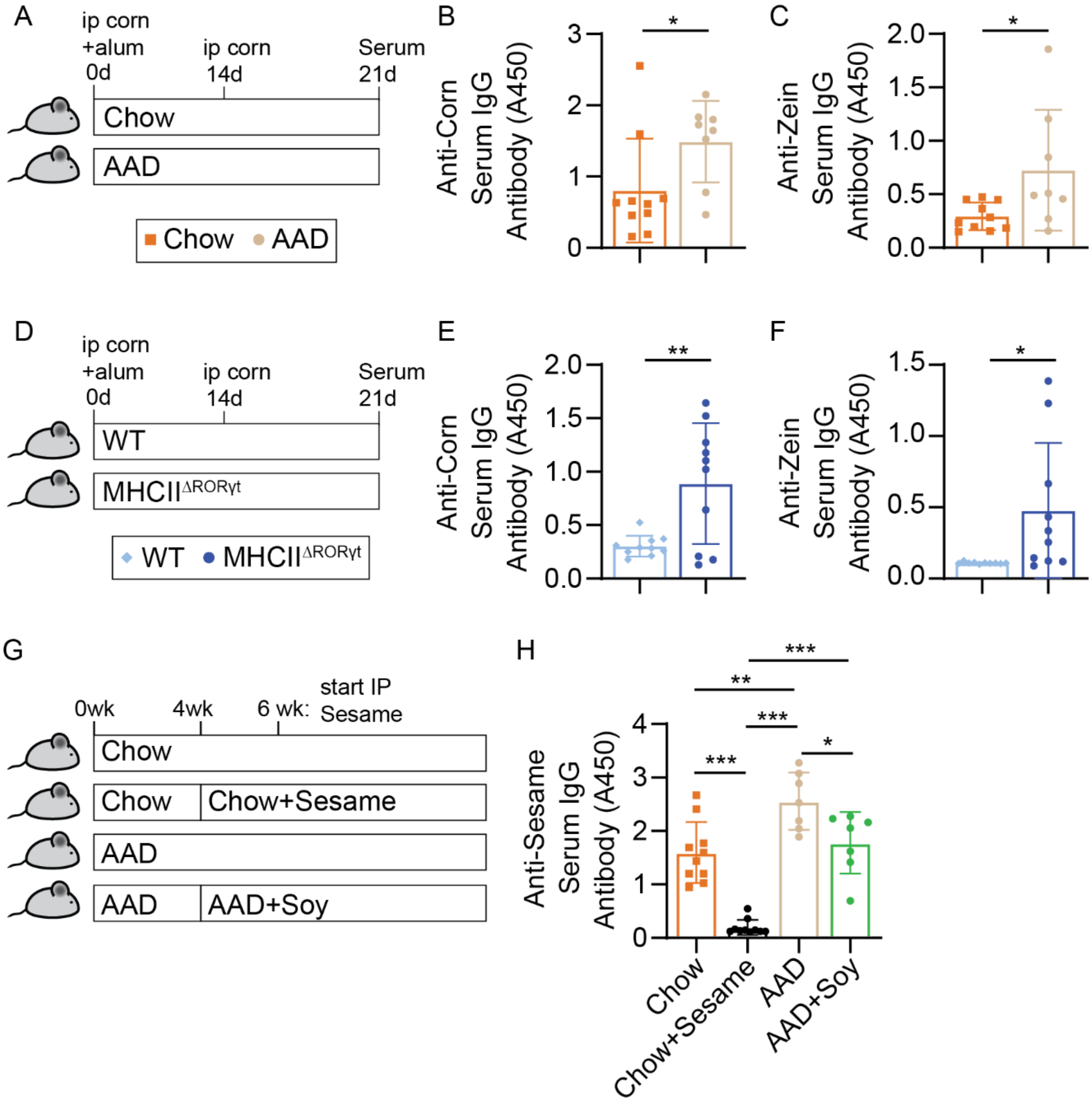


**Fig. S22. Antibody response to an inflammatory corn injection.**

**(A)** Mice born onto chow or AAD diets received an inflammatory corn exposure involving IP injection with corn+alum on day 0, a booster with IP corn on day 14 and serum collection on day 21. **(B-C)** Anti-corn and anti-zein IgG antibodies in the serum of mice on different diets injected with intraperitoneal corn in alum and boosted with corn alone. **(D)** Control mice (WT) or mice with an RORγt- driven MHCII deletion (MHCII$^{\Delta ROR\gamma t}$) on chow diet received an inflammatory corn exposure involving IP injection with corn+alum on day 0, a booster with IP corn on day 14 and serum collection on day 21. **(E-F)** Anti-corn and anti-zein IgG antibodies in the serum of mice with different genetic backgrounds injected with intraperitoneal corn in alum and boosted with corn alone. **(G)** Mice were born onto chow or AAD diets. At 4 weeks of age some chow fed mice were randomized to chow+10% sesame and some AAD fed mice were randomized to AAD+10% soy. At 6 weeks of age mice began an inflammatory sesame exposure involving IP injection with sesame + alum on day 0, a booster with IP sesame on day 14 and serum collection on day 21. **(H)** Anti-sesame IgG antibodies in the serum of mice injected with intraperitoneal sesame in alum and boosted with sesame alone. N=7-12/group. Data in panels B-C are representative of two independent experiments and data in panels E-H were generated in a singel experiment. *P* values calculated by unpaired t-test (Panels B,C,E,F) or one-factor ANOVA with a Tukey's multiple comparisons test (Panels H) Error bars indicate mean ± SD. Every dot represents an individual mouse. * denotes $p<0.05$, ** denotes $p<0.01$ and *** denotes $p<0.001$.

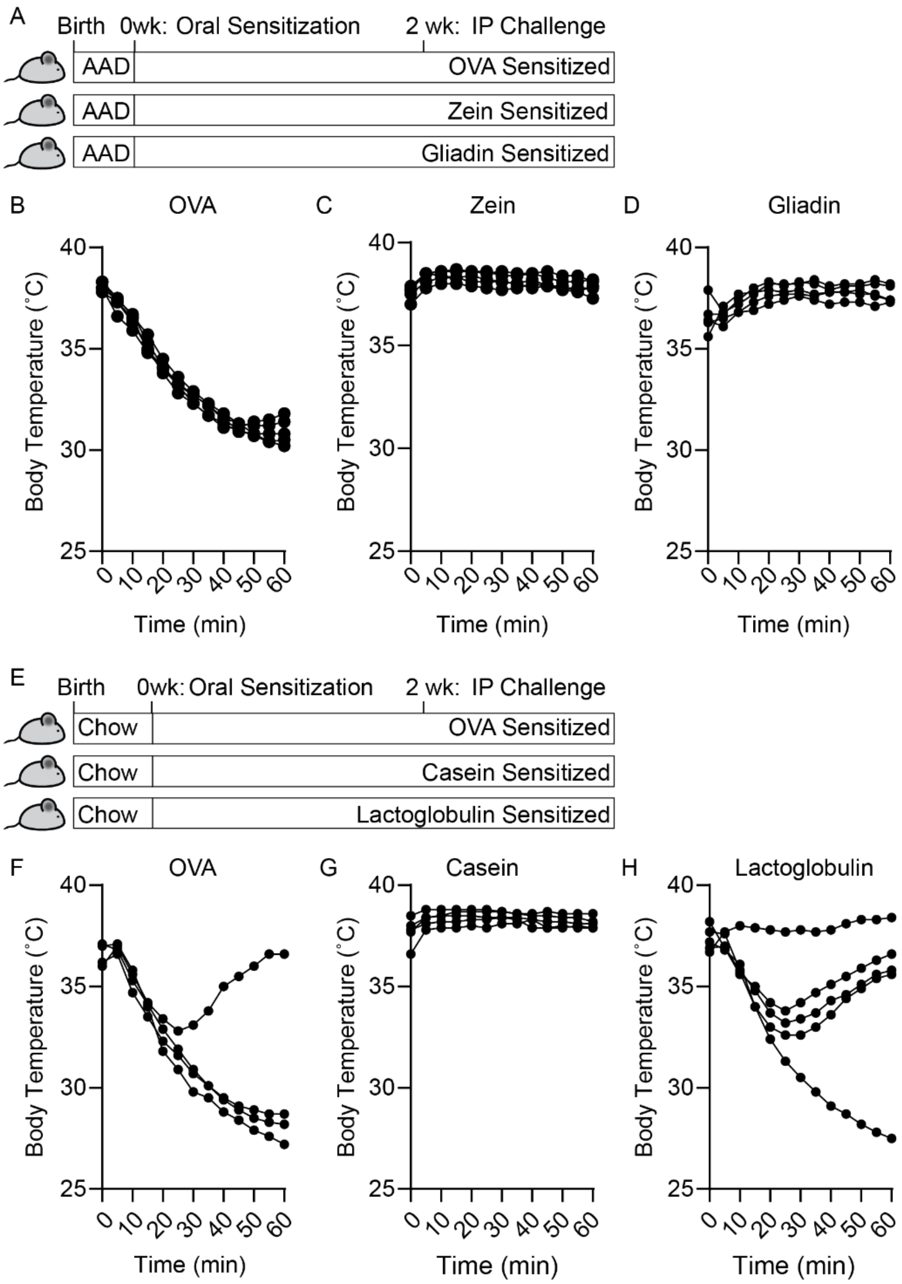


**Fig. S23. Response to various antigens in an oral allergy model**

(A) To indue an allergy, mice born onto amino acid defined diets were sensitized twice with 5 mg of protein (OVA, Zein, or gliadin) and 10 ug cholera toxin (days 0 and 7). A serum sample was collected for ELISA analysis on day 13. On day 14, mice were injected with 2 mg of their allergen and body temperature was monitored for 1 hour. **(B-D)** Body temperature following intraperitoneal challenge with OVA, Zein, or Gliadin. **(E)** To indue an allergy, mice born onto chow diets were sensitized twice with 5 mg of protein (OVA, casein, or lactoglobulin) and 10 ug cholera toxin (days 0 and 7). **(F-H)** Body temperature following intraperitoneal challenge with OVA, casein, or lactoglobulin. N=5/group in Panels B-E. Data in panels B, D, G and H were generated in a single experiment and data in panels C and F are representative of two independent experiments. *P* values calculated by unpaired t-test (Panels D-E). Error bars indicate mean ± SD. Every dot represents an individual mouse. * denotes $p<0.05$, ** denotes $p<0.01$ and *** denotes $p<0.001$

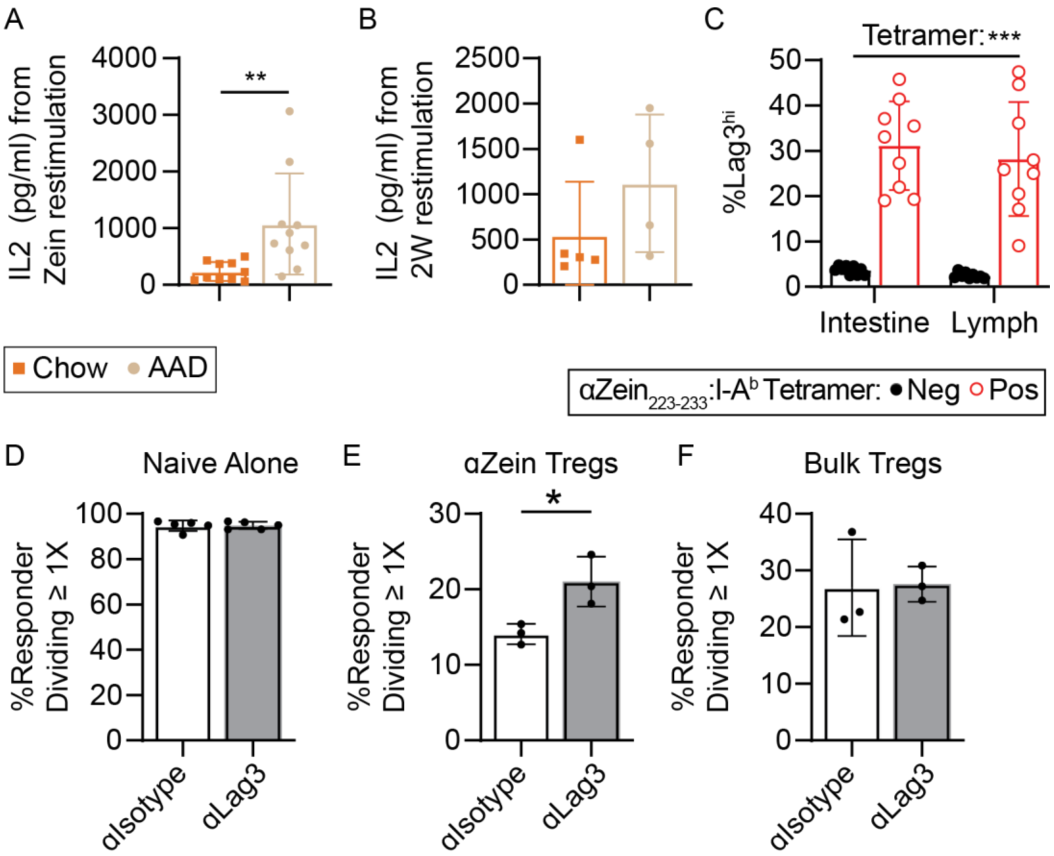


**Fig. S24. T cell profile following CFA injection for various epitopes**
**(A-B)** 8 days after epitope + CFA injection, lymph node cells were isolated and then restimulated in culture with cognate peptide for 24 hours. Media IL2 levels were measured as an indicator of T cell activation. **(C)** Lag3 expression between αZein$_{223\text{-}233}$ binding and tetramer non-binding Tregs isolated from intestine or draining lymph node following epitope + CFA injection. **(D-F)** Percent of naïve T cells divided after incubation with antigen presenting cells, a Treg population (αZein-specific Tregs or tetramer negative Tregs isolated from the inguinal lymph node after CFA peptide injection), αCD3 antibodies, and isotype targeting or CD3 targeting antibodies to selectively target Lag3 mediated suppression. N=10/group in Panel A. N=4-5/group in Panel B. N=9/group in Panel C. N=5/group in Panel D. N=3/group in Panels E-F. Data were generated in a single experiment. *P* values calculated by unpaired t-test (Panels A,B,D,E,F) or two-factor repeated measures ANOVA (Panel C). Error bars indicate mean ± SD. In Panels A-C every dot represents an individual mouse. Panels D-F represent experimental replicates from cells pooled across mice. CFA, Complete Freund's Adjuvant. * denotes $p<0.05$, ** denotes $p<0.01$ and *** denotes $p<0.001$.

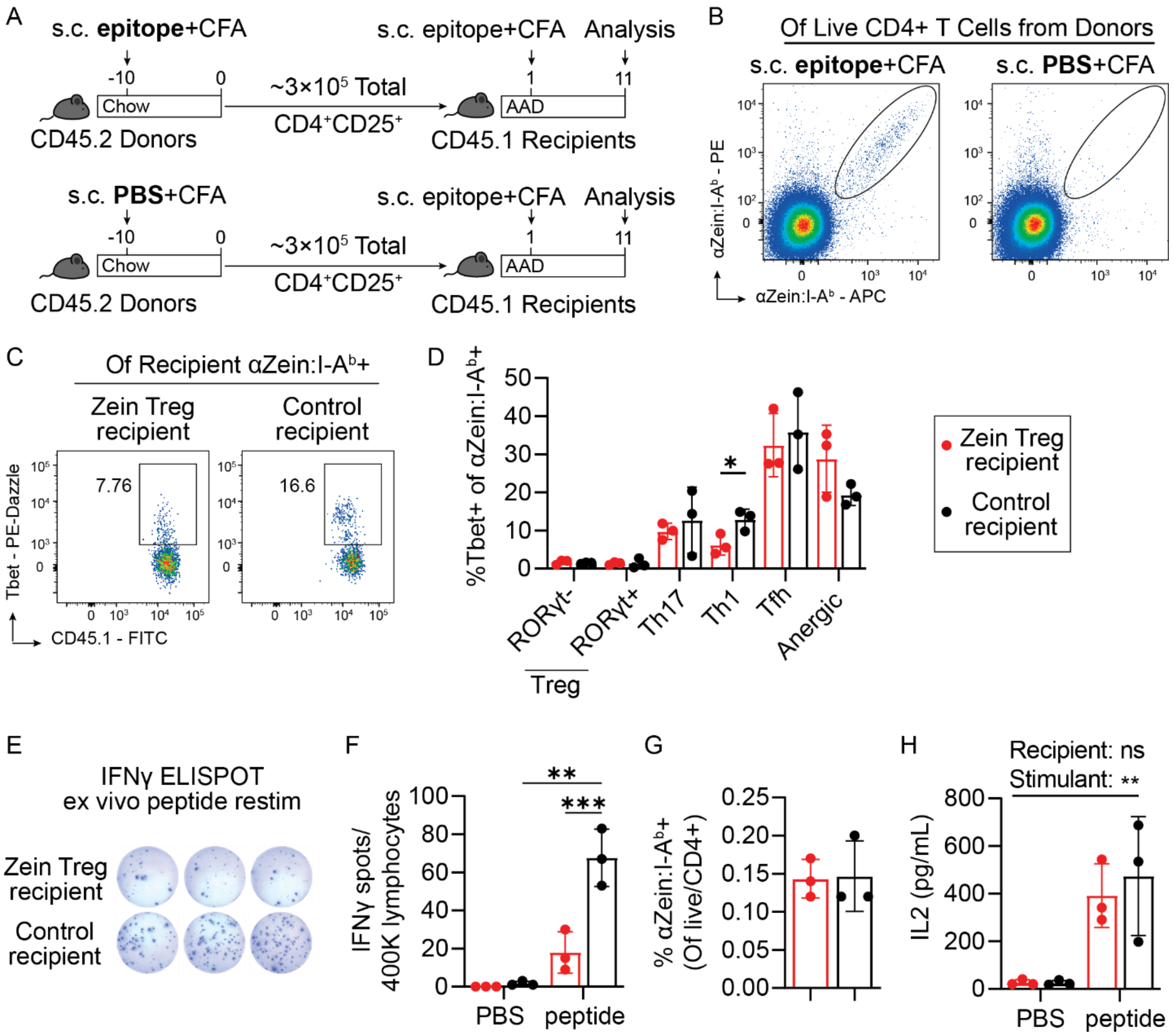


**Fig. S25. Adoptive Transfer of Zein Tregs confers protection from an inflammatory challenge. (A)** CD45.2 mice consuming chow diet were injected with either zein epitope or PBS emulsified in CFA. After 10 days total CD4+Cd25+ Tregs were sorted from the draining lymph node and injected into CD45.1 recipient mice consuming AAD diet. The following day the recipient mice were all injected with zein epitope emulsified in CFA. All analysis is of draining lymph node cells in the recipient mice. **(B)** Confirmation of Zein T cells in the donor mice. **(C-D)** Representative Tbet staining and overall distribution of cell types responsive to αZein in recipient mice. **(E-F)** Th1 cells in recipient mice identified by ELISpot assay. **(G)** Frequency of zein T cells in donor mice. **(H)** After epitope + CFA injection, lymph node cells were isolated and then restimulated in culture with zein peptide for 24 hours. Media IL2 levels were measured as an indicator of T cell activation. N=3/group. All data generated from a single experiment. *P* values were calculated using an unpaired t-test (Panels D,G), two-factor repeated measures ANOVA with a Uncorrected Fisher's LSD test (Panel F), or two-factor repeated measures ANOVA (Panel H). Complete Freund's Adjuvant. * denotes p<0.05, ** denotes p<0.01 and *** denotes p<0.001.

**Table S1:** Approaches used to select TCRs for hybridoma generation

| **Approach** | **# of TCRs selected to screen** | **# of TCRs from 1+ Treg** | **# Food responsive** | **Food-responsive TCR Name** |
|---|---|---|---|---|
| Expanded Clonotypes[1] | 55 | 32 | 3 | α_Zein_TCR_1; α_Zein_TCR_2; Gly_TCR_1 |
| Predicted microbiome signature[1,2] | 37 | 14 | 1 | α_Zein_TCR_3 |
| Expanded clonotypes from germ-free mice with at least 1 Treg | 21 | 21 | 2 | α_Zein_TCR_4; Unmapped TCR |
| In a GLIPH cluster with original 4 food responsive TCRs | 7 | 5 | 1 | α_Zein_TCR_5 |
| Preliminary gene-signature guided[3] | 4 | 4 | 1 | Unmapped TCR |
| Random selection from colonized small intestine | 4 | 4 | 1 | Gli_TCR_1 |
| Random selection from Tregs in scRNA-seq data collected in this manuscript | 5 | 5 | 0 | - |
| Selection of Tregs from Cluster 1 in scRNA-seq data collected in this manuscript | 8 | 8 | 1 | Gli_TCR_2 |

[1]These TCRs from Nagashima et al., 2023
[2]Using criteria from Nagashima et al., 2023
[3]Criteria: Is a Treg (Leiden clusters high on Foxp3/IL10); high on Klrg1 and Fam46a and low on Got1 compared to other Tregs [differential markers between food-responsive Tregs and bacterial responsive-Tregs from data in Nagashima et al., 2023]

**Table S2:** Antibodies used in experiments

| Antibody | Clone | Vendor; RRID | Catalog Number | Dilution |
|---|---|---|---|---|
| eBioscience Fixable Viability Dye eFluor 780 | | Thermo Fisher | 65-0865-14 | 1 in 750 |
| Brilliant Ultra Violet 496 anti-mouse CD4 | GK1.5 | BD Bio; AB_2813886 | 612952 | 1 in 200 |
| Brilliant Violet 711 anti-mouse CD4 | RM4-5 | Biolegend; AB_11219396 | 100549 | 1 in 200 |
| Brilliant Violet 510 anti-mouse CD4 | RM4-5 | Biolegend; AB_2561388 | 100553 | 1 in 200 |
| PE/Cyanine7 anti mouse CD49b | HMα2 | Biolegend; AB_2566102 | 103517 | 1 in 200 |
| APC anti-mouse LAG3 | C9B7W | Biolegend; AB_10639935 | 125209 | 1 in 200 |
| PerCP/Cyanine5.5 anti-mouse FR4 (Folate Receptor 4) | 12A5 | Biolegend; AB_2721723 | 125017 | 1 in 200 |
| Brilliant Violet 605 anti-mouse CD73 | TY/11.8 | Biolegend; AB_2561528 | 127215 | 1 in 200 |
| PE Foxp3 | FJK-16s | Thermo Fisher; AB_465936 | 12-5773-82 | 1 in 100 |
| FITC anti-human/mouse Helios | 22F6 | Biolegend; AB_10549181 | 137204 | 1 in 200 |
| Brilliant Violet 650 C25 | PC61.5 | Thermo Fisher; AB_468914 | 16-0251-85 | 1 in 200 |
| Brilliant Violet 421 anti-mouse LAP | TW7-16B4 | Biolegend; AB_2561580 | 141407 | 1 in 200 |
| APC Foxp3 | FJK-16s | Thermo Fisher; AB_469457 | 17-5773-82 | 1 in 100 |
| PE/Cyanine7 anti-mouse Foxp3 | FJK-16s | Thermo Fisher; AB_891552 | 25-5773-82 | 1 in 100 |
| Brilliant Violet 711 anti-mouse GATA3 | L50-823 | BD Bio; AB_2739242 | 565449 | 1 in 100 |
| PE-CF594 anti-mouse Tbet | Apr-46 | BD Bio; AB_2737621 | 562467 | 1 in 100 |
| Brilliant Violet 421 anti-mouse RORγt | Q31-378 | BD Bio; AB_2687545 | 562894 | 1 in 100 |
| Brilliant violet 510 anti-mouse CD4 | RM4-5 | Biolegend; AB_2561388 | 100553 | 1 in 200 |
| Pe/Dazzle 594 anti-mouse CD223 (Lag-3) | C9B7W | Biolegend; AB_2572081 | 125223 | 1 in 200 |

| | | | | |
|---|---|---|---|---|
| PE/Cyanine7 anti-human/mouse Granzyme B | QA16A02 | Biolegend; AB_2728380 | 372213 | 1 in 200 |
| Brilliant Violet 510 anti-mouse CD279 (PD-1) | 29F.1A12 | Biolegend; AB_2715761 | 135241 | 1 in 200 |
| PerCP/Cyanine5.5 anti-mouse CD152 | UC10-4B9 | Biolegend; AB_2564473 | 106315 | 1 in 200 |
| FITC anti-mouse CD45.1 | A20 | Biolegend; AB_313494 | 110705 | 1 in 200 |
| eFluor 450 anti-mouse CD45R | RA3-6B2 | Thermo Fisher; AB_1548761 | 48-0452-82 | 1 in 200 |
| Brilliant Violet 510 anti-mouse TCRgd | GL3 | Biolegend; AB_2563534 | 118131 | 1 in 200 |
| Brilliant Violet 650 anti-mouse CD45.1 | A20 | BD Bio; AB_2738405 | 563754 | 1 in 200 |
| Brilliant Violet 711 anti-mouse TCRb | H57-597 | BD Bio; AB_2738023 | 563135 | 1 in 200 |
| PerCP/Cyanine5.5 anti-mouse CD45.2 | 104 | Biolegend; AB_893350 | 109828 | 1 in 200 |
| AlexaFluor 700 anti-mouse CD45.2 | 104 | Biolegend; AB_493730 | 109821 | 1 in 200 |
| APC anti-mouse TCRvb6 | RR4-7 | Thermo Fisher; AB_11041976 | 17-5795-82 | 1 in 200 |
| Alexa Fluor 700 anti-mouse CD4 | RM4-5 | Thermo Fisher; AB_494000 | 56-0042-82 | 1 in 200 |
| APC anti-mouse CD62L | MEL-14 | Thermo Fisher; AB_469411 | 17-0621-83 | 1 in 200 |
| PerCP/Cyanine 5.5 anti-mouse CD44 | IM7 | Thermo Fisher; AB_925746 | 45-0441-82 | 1 in 200 |
| FITC anti-mouse TCRvb6 | RR4-7 | BD Bio; AB_394700 | 553193 | 1 in 200 |
| PE anti-mouse CD25 | PC61.5 | Thermo Fisher; AB_465607 | 12-0251-82 | 1 in 200 |
| FITC anti-mouse CD4 | GK1.5 | Cytek; AB_2621665 | 35-0041 | 1 in 200 |
| PerCP/Cyanine5.5 anti-mouse MHC class II (I-A/I-E) | M5/114 | BD Bio; AB_11153297 | 562363 | 1 in 200 |
| PerCP/Cyanine5.5 anti-mouse CD11b | M1/70 | Thermo Fisher; AB_953558 | 45-0112-82 | 1 in 200 |

| | | | | |
|---|---|---|---|---|
| PerCP/Cyanine5.5 anti-mouse CD11c | N418 | Thermo Fisher; AB_925727 | 45-0114-82 | 1 in 200 |
| PerCP/Cyanine5.5 anti-mouse B220 | RA3-6B2 | Thermo Fisher; AB_1107006 | 45-0452-82 | 1 in 200 |
| PE-Dazzle anti-mouse CD44 | IM7 | Biolegend; AB_2564043 | 103055 | 1 in 200 |
| APC anti-mouse CD62L | MEL-14 | Biolegend | 104411 | 1 in 200 |
| PE anti-mouse CX3CR1 | SA011F11 | Biolegend; AB_2564314 | 149005 | 1 in 200 |
| Brilliant Violet 421 anti-mouse MHCII | M5/114.15.2 | Biolegend; AB_10900075 | 107631 | 1 in 200 |
| Alexa Fluor 700 anti-mouse TCRβ | H57-597 | Biolegend; AB_1027654 | 109223 | 1 in 200 |
| Brilliant Violet 711 anti-mouse F4/80 | BM8 | Biolegend; AB_2564588 | 123147 | 1 in 200 |
| Brilliant Violet 650 anti-mouse CD11c | N418 | Biolegend; AB_2562414 | 117339 | 1 in 200 |
| FITC anti-mouse TCRβ | H57-597 | Biolegend; AB_313428 | 109205 | 1 in 200 |
| Brilliant Violet 711 anti-mouse CD366 (Tim-3) | RMT3-23 | Biolegend; AB_2716208 | 119727 | 1 in 200 |
| Brilliant Violet 510 anti-mouse PD1 | 29F.1A12 | Biolegend; AB_2715761 | 135241 | 1 in 200 |
| PE anti-mouse IL10 | JES5-16E3 | Biolegend; AB_315361 | 505007 | 1 in 200 |
| Brilliant Violet 785 anti-mouse/human Klrg1 | 2F1/KLRG1 | Biolegend; AB_2629749 | 138429 | 1 in 200 |
| APC anti-mouse CD366 (Tim-3) | B8.2C12 | Biolegend; AB_2562997 | 134007 | 1 in 200 |
| PE/Cyanine7 anti-mouse CD45.1 | A20 | Thermo Fisher; AB_469629 | 25-0453-82 | 1 in 200 |
| Brilliant Violet 650 anti-mouse CD8a | 53-6.7 | Biolegend; AB_11124344 | 100741 | 1 in 200 |

**Data file S1. (separate file)**
S1: Plants sharing an exact epitope with $Gliadin_{273-285}$.

**Data file S2. (separate file)**
S2: Top candidate proteins for cross-reactivity identified through BLASTP analysis for Gly_TCR_1.

**Data file S3. (separate file)**
S3: Top candidate proteins for cross-reactivity identified through BLASTP analysis for αZein_TCRs.

**Data file S4. (separate file)**
S4: Differentially expressed genes identified from bulk RNA-sequencing

**Data file S5. (separate file)**
S5: TCR Sequences on food-reactive hybridomas.

**Data file S6. (separate file)**
S6: Tabulated data for Fig. 1-5, S1-S25.
